# Understanding the Free Energy Landscape of Phase Separation in Lipid Bilayers using Molecular Dynamics

**DOI:** 10.1101/2023.01.31.526537

**Authors:** Ashlin J. Poruthoor, Akshara Sharma, Alan Grossfield

## Abstract

Liquid-liquid phase separation (LLPS) inside the cell often results in biological condensates that can critically impact cell homeostasis. Such phase separation events occur in multiple parts of cells, including the cell membranes, where the so-called “lipid raft” hypothesis posits the formation of ordered domains floating in a sea of disordered lipids. The resulting lipid domains often have functional roles. However, the thermodynamics of lipid phase separation and their resulting mechanistic effects on cell function and dysfunction are poorly understood. Understanding such complex phenomena in cell membranes, with their diverse lipid compositions, is exceptionally difficult. For this reasons, simple model systems that can recapitulate similar behavior are widely used to study this phenomenon. Despite these simplifications, the timescale and and length scales of domain formation pose a challenge for molecular dynamics (MD) simulations. Thus, most MD studies focus on spontaneous lipid phase separation — essentially measuring the sign (but not the amplitude) of the free energy change upon separation — rather than directly interrogating the thermodynamics. Here, we propose a proof-of-concept pipeline that can directly measure this free energy by combining coarse-grained MD with enhanced sampling protocols using a novel collective variable. This approach will be a useful tool to help connect the thermodynamics of phase separation with the mechanistic insights already available from molecular dynamics simulations.

**SIGNIFICANCE:** Standard molecular dynamics simulations can determine the sign the free energy change upon phase separation, but not the amplitude. We present a new method to determine the phase separation free energy for lipid membranes, based on a enhanced sampling using the weighted ensemble method combined with a novel collective variable, validated using coarse-grained simulations applied to several simple systems. The new method will be valuable as a way to develop models that connect molecular-level structural features to the thermodynamics of phase separation.

## INTRODUCTION

Interactions among biomolecules often result in phase separation and subsequent formation of biological condensates(1). In the past decade, there has been a new appreciation for the role of phase separation in cell physiology(2). Biological condensates involving fundamental biomolecules like DNA(3), RNA(4), protein(5), and lipids(6) have been identified and their functional roles at least somewhat understood. Biological condensates are involved in diverse processes, including DNA damage response(7), translational(8) and transcriptional(9) regulation, ribosome biogenesis(10), cell adhesion(11), and endocytosis(12). Such nano-to-micron-scale compartments have no surrounding membrane but sequester key molecules and help in cellular organization, similar to canonical organelles(13, 14). These transient and dynamic sequestration zones are crucial for cell stress responses(14) and signal transductions(15). As phase-separated molecules are concentrated within these condensates, they can function as reaction crucibles that enhance reaction kinetics(16). However, abnormal phase behavior of biological condensates is speculated to underlie multiple disease states, including neurodegenerative diseases such as Huntington’s(17), ALS(18), and Parkinson’s(19).

Like these intracellular biological condensates, phase separation in the cell membrane often results in relatively ordered lipid lateral domains with collective behavior that can recruit other proteins and lipids(6, 20). Such domains often cluster signaling molecules(21) and facilitate conformational modulations through domain-specific lipid interactions(22, 23) that are relevant in immune signaling(24, 25) and host-pathogen interactions(26). As membrane properties modulate resident protein function, the coexistence of distinct phases gives the cell an extra tool for regulation. Conformational changes in domain-resident molecules and their subsequent activity shifts regulate key signaling events(22, 23). It has also been observed that the HIV Gag protein is sensitive to domains with high cholesterol content, suggesting a potential role of membrane domains in host-pathogen interaction and viral assembly(26). Moreover, lipid domains are conserved throughout the tree of life, implying their relevance in regulating cellular processes(6); for example, recent work from the Keller lab showed that yeast actively regulate the composition of their vacuolar membranes to keep the melting temperature well above the growth temperature of the yeast(27). For these reasons, understanding the thermodynamics of lipid phase separation as a function of the composition of the system is a critical step toward understanding the individual lipid properties that underlie the formation of domains and the resultant functional modulations.

Lipid domains have been studied extensively both experimentally and computationally. However, teasing out the complex interactions between domain components that determine the membrane organization is challenging in vivo due to the limitations of different methodologies(28). Such difficulties primarily arise due to (a) the complex composition of cell membrane(29), (b) difficulties in defining domain properties in vivo(6), and (c) challenges in achieving specificity when perturbing the system with probes(30). Hence, simple model membranes have been extensively used to mimic the phase separation of complex cell membrane in vitro(31–33) and in silico(34–36). Such studies provide powerful insights into phase-separated domains, their properties, and whether the process is favorable, but as a rule atomistic and coarse-grained models cannot quantify the underlying thermodynamics, although simpler lattice-based models can (37–45). This limits our ability to assess the contributions of different mechanistic effects to the process.

As a “computational microscope(46)”, molecular dynamics (MD) simulations have been used to study spontaneous lipid separation without using exogenous labels or probes(47). However, MD simulations designed for spontaneous phase separation are inadequate to compute thermodynamic properties with statistical confidence. This is because the transition between mixed and separated states is often a single irreversible event on the microsecond timescale typically achievable by typical all-atom or even coarse-grained MD(34, 48), especially given the relatively large size required to model the phenomenon. That said, CG-MD is an excellent tool to study phase separation in the membrane, because it retains molecular-level resolution while capturing nanoscale domains due to transient phase separation events.

Previously, the Tieleman(48) and the Gorfe groups(36) have done some pioneering work on the thermodynamics of lipid domains. While the former used thermodynamic integration(49) to calculate the free energy and excess chemical potential for exchanging lipid species between phases, the latter used umbrella sampling(50) to compute the membrane partition thermodynamics of an idealized transmembrane domain by dragging it across the boundary between distinct lipid phases.

We hypothesize that coupling a more versatile enhanced sampling method with the standard CG-MD will improve the transition events between the mixed and separated states of the lipid bilayer system under study to compute the free energy for phase separation. Various enhanced sampling protocols have previously demonstrated their ability to enhance the sampling of rare events(51). In most cases, external forces are applied to the system to bias simulation to explore the desired phase space. As a result, it is necessary to compute the derivative of the collective variable at each time step. Certain popular implementations of such protocols, like COLVARS(52) and PLUMED(53), are done on single CPUs serially, leading to poor performance when the collective variable is computationally complex, as is the case for the complicated collective variables proposed to study phase separation.

By contrast, the weighted ensemble (WE)(54, 55) method has the advantages of (a) unbiased dynamics, and (b) good sampling, achieved by enhancing the sampling of otherwise undersampled phase space and reducing the sampling of otherwise oversampled phase space. Moreover, there is an added benefit, in that the calculation of the collective variable can be decoupled from the MD, which simplifies its implementation and can improve the computational performance. It should be noted that the WE method is highly parallelizable and can take advantage of GPU acceleration(56).

Here we present a novel proof-of-concept pipeline to estimate the equilibrium free energy change upon separating a lipid bilayer into distinct coexisting phases. We test this pipeline on three different lipid bilayer systems with varying degrees of phase separation propensity. We explore three potential candidate collective variables and assess their effectiveness in the pipeline. We then construct free energy landscapes of the three systems at multiple temperatures. We further explore a few non-traditional use cases for the data thus generated and discuss additional room for improvements.

## METHODS

### System details

As shown in Fig 1, we used three different ternary lipid bilayer systems: 1. A lipid bilayer consisting of dipalmitoyl-phosphatidylcholine (DPPC), dilinoleyl-phosphatidylcholine (DIPC), and cholesterol (CHOL), known to phase separate in silico in a few microseconds (15, 34, 59–63). 2. A lipid bilayer consisting of DPPC, diarachidonoyl-phosphatidylcholine (DAPC), and CHOL, known to phase separate in silico within a few hundred nanoseconds (35, 36, 64). 3. A lipid bilayer consisting of DPPC, palmitoyl-oleoyl-phosphatidylcholine (POPC), and CHOL that was previously shown not to phase separate (33, 64). The composition of the DPPC-DIPC-CHOL, DPPC-DAPC-CHOL, and DPPC-POPC-CHOL systems used here are (0.42/0.28/0.3), (0.5/0.3/0.2) and (0.4/0.4/0.2) respectively and are adapted from previous studies (34, 35, 64). We kept the total number of lipids the same as in previous calculations as well, with the DPPC-DIPC-CHOL, DPPC-DAPC-CHOL, and DPPC-POPC-CHOL systems containing 1944, 1200 and 1200 respectively.

**Figure 1:**
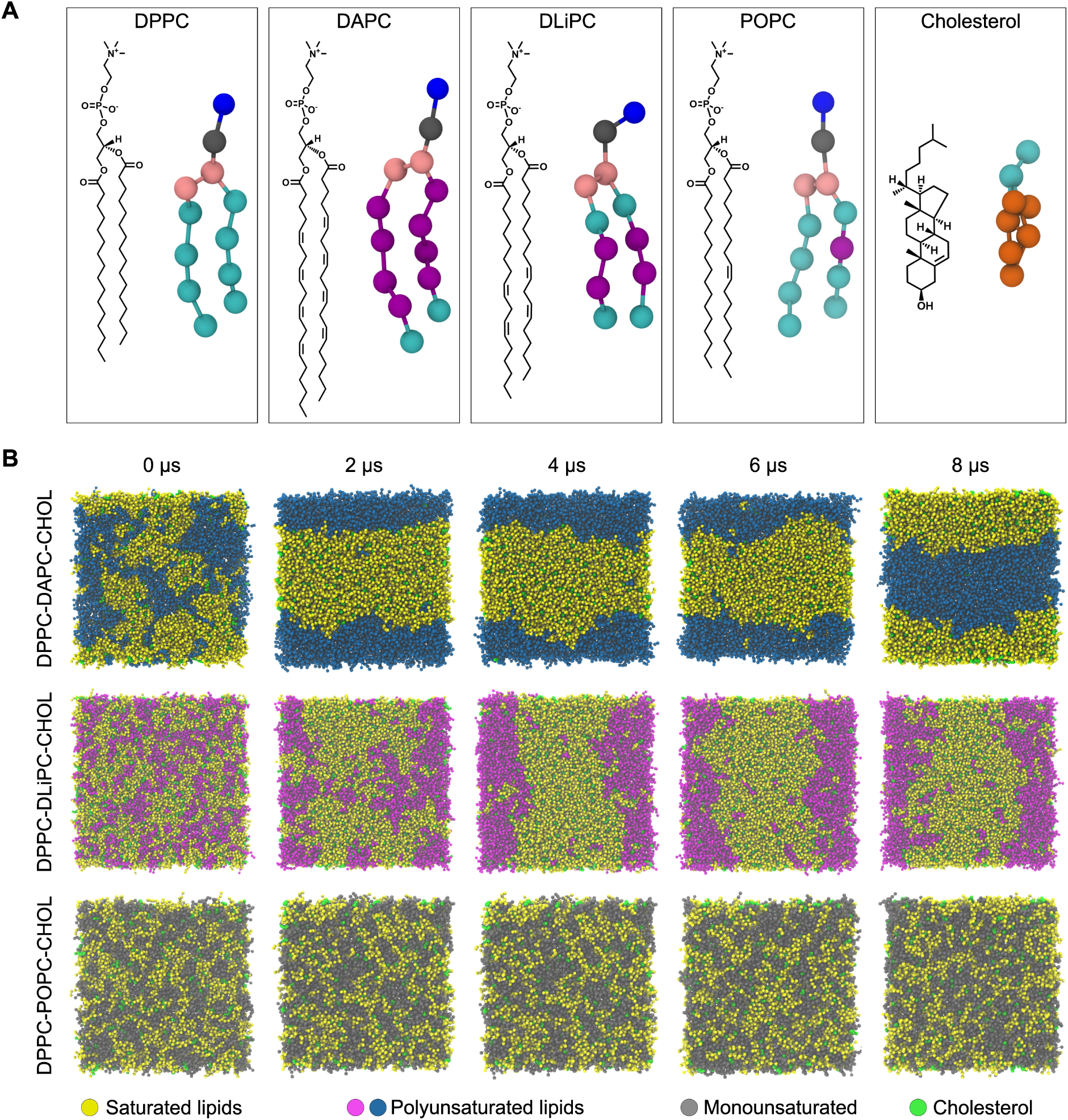
A. 2D structure and corresponding MARTINI 2 bead structure for lipids used in this study (Visualization using ChemDraw and VMD(57, 58) respectively). B. Time evolution of each lipid system as a function of time. The membrane normal is pointing towards the reader.

Due to the relatively large system sizes and the time scale required for phase separation and related dynamics in lipid bilayer simulations, we used a coarse-grained (CG) model for each system. Although nothing in the subsequent sampling or analysis is specific to the coarse-grained models, it makes sense to use a less expensive model while working out the method.

Using CHARMM-GUI Martini Maker (65), we constructed four replicas of each CG ternary symmetric bilayer system, with the lipids randomly mixed in the bilayer. We used MARTINI 2 force field parameters and particle definitions(66, 67) to construct CG systems and to run the subsequent MD simulation. We replaced the default input files from CHARMM-GUI Martini Maker with their respective most recent Martini 2.x versions if they existed. We used the MARTINI polarizable water model(68) to solvate all systems with approximately a 1:30 lipid to real water ratio. A detailed description of the systems is given in Table S1 of supplementary material. We did not use the more recent MARTINI 3.0 lipid parameters, since there are no published MARTINI 3 parameters for cholesterol at this time.

### Standard MD simulation details

We used GROMACS 2020.3(69) to propagate the dynamics of the systems prepared. Each system was minimized and equilibrated following the protocol suggested by CHARMM-GUI Martini Maker. To obtain an intact bilayer without any membrane undulations, we used the flat bottomed restraint potential available in GROMACS; this allowed the lipids to move freely in the plane of the membrane but restrained within a slab of defined 𝑧 thickness along the membrane normal. More details about membrane restraining protocol are given in the supplementary material (section 2).

After the minimization and equilibration, all systems were run at 400 K in the NPT ensemble for 100 ns to make sure the lipids in each system were randomly distributed. For every system, we forked each replica into multiple temperature runs simulated at different temperatures ranging from 298K to 450K. For these production runs, the temperature coupling is done using velocity rescaling(70) with a time constant 1 ps. An extended-ensemble Parrinello-Rahman pressure coupling(71) with a relatively high time constant of 12 ps was used, as recommended for MARTINI. A semi-isotropic pressure coupling suited for membrane simulations is used here with a compressibility of 3×10^−4^ bar^−1^ and reference pressure for coupling as 1 bar. Reaction field electrostatics with a Coloumb cutoff of 1.1 nm and a dielectric constant of 2.5 was used, as required with the MARTINI polarizable water model. For Van der Waals interaction, a similar cutoff of 1.1 nm was used. A potential-shift Van der Waal modifier was also used. For neighbor searching, a verlet cutoff scheme was used with neighbor list updated every 20 steps. The simulation parameters were mostly inspired by previous CG MARTINI simulations(72). To eliminate potential artifacts previously reported due to inaccurate constraints(73), we used an 8th order expansion for the LINCSolver constraint coupling matrix(74) for more accuracy. Moreover, for better energy conservation, we also used short 20 fs timesteps, which is conservative for a CG system.

All standard MD simulations ran for at least 8 microseconds using the BlueHive supercomputing cluster of the Center for Integrated Research and Computing at the University of Rochester. Simulations ran on Intel Xeon E5-2695 and Gold 6130 processors augmented with Tesla K20Xm, K80, and V100 GPUs. The trajectories were processed and analyzed using the LOOS software package(75). A detailed description of the simulation parameters is given in Table S1 of supplementary material.

### Collective Variables

A collective variable is a reduced coordinate that captures the progress of a system along the transition of interest. Ideally, such a reduced variable(s) should fully capture the key modes of the system to reflect the complex event under study. The success and efficiency of any enhanced sampling protocol depends on the chosen collective variable over which the sampling is enhanced(51, 76, 77). We explored 3 candidate collective variables for the WE simulations.

#### 1. Fraction of Lipids in Clusters (FLC)

Since the formation of lipid domains with distinct properties from the rest of the bilayer is a characteristic feature of a phase-separating lipid bilayer, we hypothesized that we could use a variable that quantifies the recruitment of lipids into such domains to track the phase separation events in our systems. Here, we define the Fraction of Lipids in Clusters (FLC) as follows:

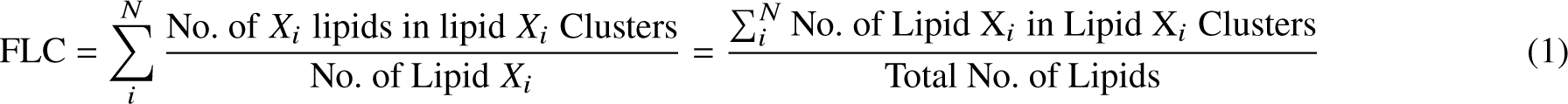

where subscript 𝑖 denotes the individual lipid species in a bilayer consisting of N total lipid species. As shown in Figure 2, each system under study has 𝑁 = 3 lipid species. FLC increases as the system goes from a well-mixed state, with a random distribution of lipids, to a separated state. An FLC of 0 corresponds to a system configuration where no lipids are part of any cluster, while an FLC of 1 implies that all lipids are in a cluster (though not necessarily a single cluster).

**Figure 2:**
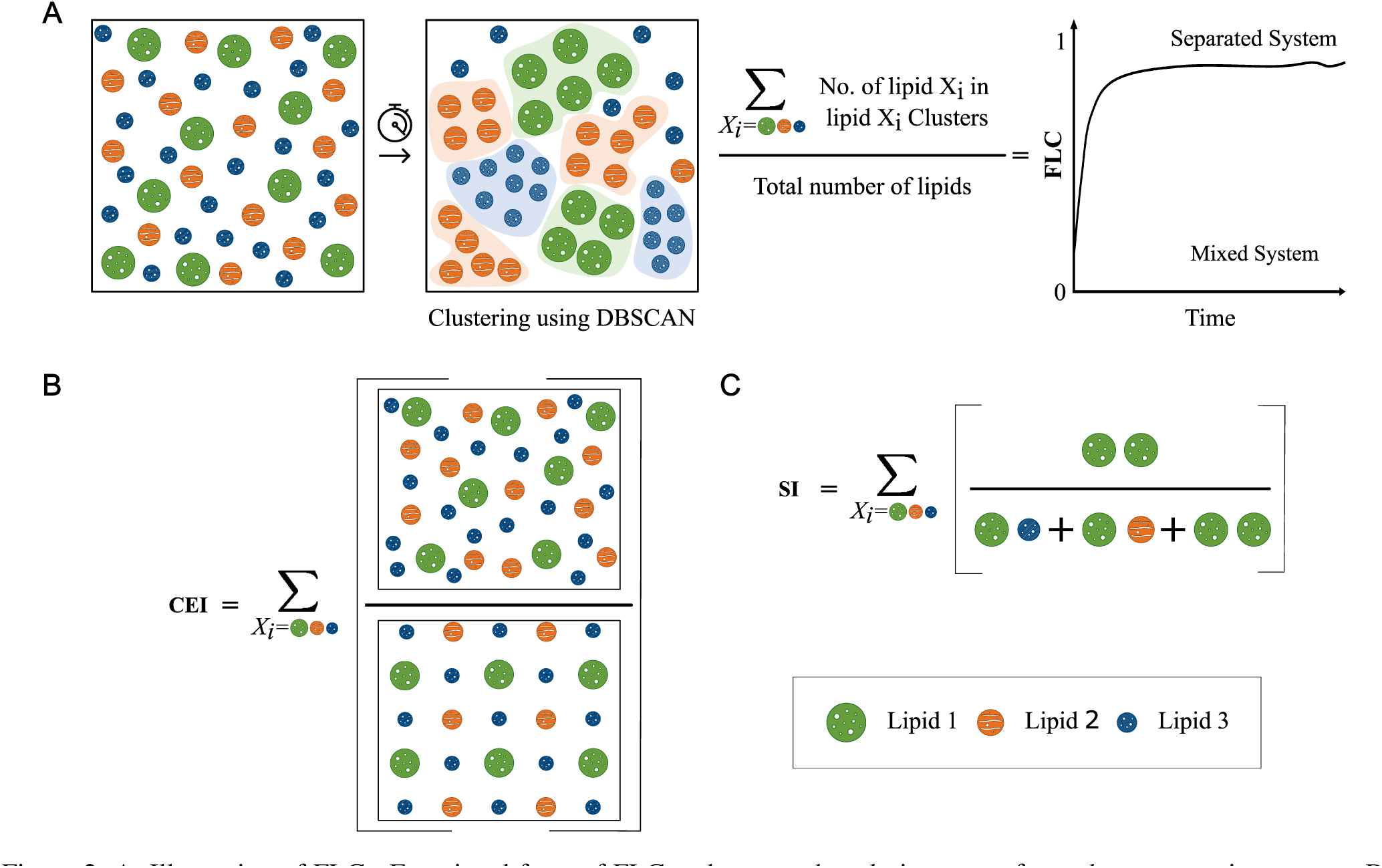
A. Illustration of FLC : Functional form of FLC and expected evolution curve for a phase separating system. B. Illustration of Cumulative Enrichment Index. C. Illustration of Segregation Index

The clustering is defined using the Density-Based Spatial Clustering of Applications with Noise (DBSCAN) algorithm (78, 79) as implemented in scikit-learn (80). For lipid DBSCAN clustering, we supply a 2-dimensional lipid-lipid distance matrix that accounts for periodic boundary conditions; the distance calculation was performed using LOOS, with the two leaflets treated independently. The DBSCAN algorithm requires two additional input parameters: 𝑚𝑖𝑛_𝑠𝑎𝑚 𝑝𝑙𝑒𝑠 and 𝜀. We consider lipids with more than 𝑚𝑖𝑛_𝑠𝑎𝑚 𝑝𝑙𝑒𝑠 neighbors (including the lipid itself) within 𝜀 radius as core lipids. Non-core lipids still within 𝜀 radius of a core lipid are considered border lipids. A set of core lipids within 𝜀 radius of each other and their border lipids forms a cluster. All lipids that are not a part of any cluster are considered outliers.

Since lipid motion in a bilayer is constrained primarily on a plane and MARTINI beads for a lipid are of similar radii, we used the two-dimensional version of Kepler’s conjecture that the densest packing of unit disks in a plane is hexagonal close packing (Thue’s Theorem), and chose 7 (6 nearest neighbors + 1 central lipid) as 𝑚𝑖𝑛_𝑠𝑎𝑚 𝑝𝑙𝑒𝑠 for all the lipid species. However, 𝜀 was chosen differently for each lipid species based on their first nearest-neighbor distance from the central lipid. We used the 𝑥𝑦_𝑟 𝑑𝑓 tool in LOOS to calculate an individual lipid species’ first nearest-neighbor distance. From the first 8 𝜇s MD standard simulation of each replica, we computed the radial distribution function (RDF) for a lipid species in the xy-plane. From the RDF plot, we found the first maxima (provided it is above 1), and the distance to the minima right after this first maximum was determined to be the first nearest neighbor distance for that lipid species. This distance was averaged over all four replicas for a given system at a given temperature and then assigned as the respective 𝜀 input. Since nearest neighbor distance is a function of temperature, for the same lipid species in the same system, 𝜀 may be different for different temperatures. The computed 𝜀, i.e., average first nearest neighbor distance for different conditions, are plotted in Supplementary Figure S1. Additionally, we tracked auxiliary variables (AVs) that evaluate the quality of DBSCAN clustering since it is critical for defining the FLC that drives the WE equilibrium dynamics (Supplementary section 4, Figures S2-S5).

#### 2. Cumulative Enrichment Index (CEI)

When lipids segregate into phases, one result is that the local concentration of a given lipid type in the vicinity of other lipids of the same type is increased. We attempted to exploit this phenomenon by tracking the degree of local lipid enrichment; this was inspired by previous work from our lab (81), although other groups used similar methods (82, 83). Here, we calculated the average local density of 𝑋_𝑖_ lipids around a single lipid 𝑋_𝑖 𝑗_, within a cutoff radius, 𝜖_𝑖_, defined as above for FLC estimation. We defined a normalization factor, Φ_𝑖_, as the local density of 𝑋_𝑖_ lipids for a uniformly well-mixed system of similar composition. The ratio of former respective to latter forms the enrichment index for a lipid species, 𝑋_𝑖_. CEI is defined as the sum of individual enrichment index for all the lipid species in the system, as follows:

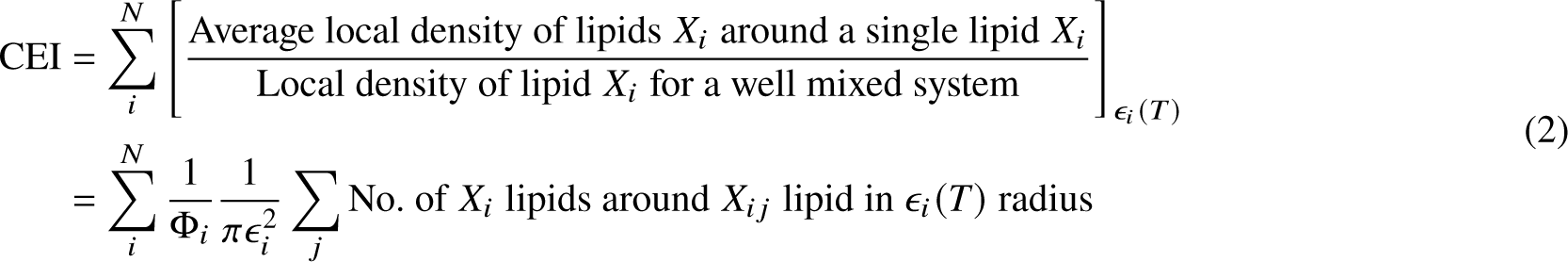

where 𝑋_𝑖 𝑗_ denotes 𝑗 ^𝑡ℎ^ lipid of 𝑋_𝑖_ lipid species. The local density around 𝑋_𝑖 𝑗_ is calculated within 𝜖_𝑖_ (𝑇) distance, where T is the temperature of the system, corrected for the contribution of the central lipid. For the normalization factor, Φ_𝑖_, we calculated the global density of 𝑋_𝑖_ lipids by taking the ratio of the total number of 𝑋_𝑖_ lipids in the system to the 𝑥𝑦-planar area of the bilayer system. This global density is the same as the local density of 𝑋_𝑖_ lipid for a uniformly well-mixed system. Thus CEI significantly larger than 3 implies that the ternary system, 𝑁 = 3, deviates from a well-mixed state to a separated state.

#### 3. Segregation Index (SI)

We defined a contact-based quantity to track the homogeneity of the lipid bilayer, similar to the ones that track the mixing of beads used previously(84, 85). Here, we computed the total contacts between a given lipid and its environment and computed the fraction of the those contacts to like-species lipids:

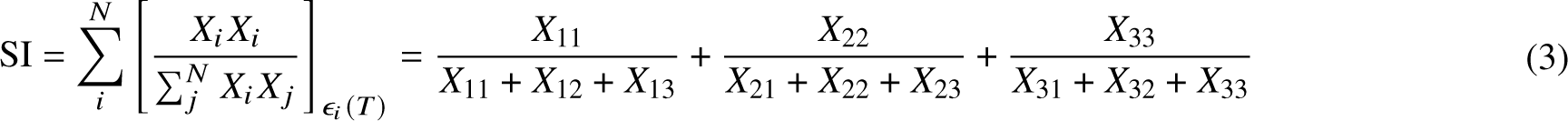

𝑋_𝑖_ 𝑋 _𝑗_ denotes the contacts between lipid species 𝑋_𝑖_ and 𝑋 _𝑗_ within 𝜖_𝑖_ (𝑇) cutoff. Thus for a ternary bilayer system, SI = 3 implies a fully separated system, and SI < 3 implies mixing. However, for the analysis, we ignored the contribution of cholesterol as we found that excluding the cholesterol term did not change the functional behavior (Supplementary Figure S6). Hence, SI_noCHOL_ effectively will have bounds [0, 2] unless otherwise stated.

### Optimized version of FLC

We used a Hidden Markov Model (HMM) (86) refinement scheme to optimize the parameters for FLC in order to improve its ability to discriminate between states. First, we calculated FLC for various combinations of 𝑚𝑖𝑛_𝑠𝑎𝑚 𝑝𝑙𝑒𝑠 and 𝜀 for the conventional MD simulations. We scanned 𝑚𝑖𝑛_𝑠𝑎𝑚 𝑝𝑙𝑒𝑠 (the number of lipids needed to form a cluster) from 5 to 60 in steps of 1 and 𝜀 from 1 nm to 𝑟_𝑑𝑖𝑎𝑔𝑜𝑛𝑎𝑙_ in an increment of 0.05 nm, where 𝑟_𝑑𝑖𝑎𝑔𝑜𝑛𝑎𝑙_ is half of the periodic box diagonal distance in the plane of the membrane. We calculated the FLC value for each parameter combination on the trajectories collected from standard MD simulation for each system at different temperatures. On the FLC evolution data corresponding to each trajectory, we fit an HMM model using the hmmlearn python module (86). However, one needs to specify the number of hidden states in the data to fit the data. Even though we could make a simple assumption of 2 states, mixed and demixed, and go ahead with fitting, it might not fully capture the behavior of the trajectory under consideration as there can be hidden meta-stable states critical to the phenomenon we are studying. Therefore, we used a brute force approach to fitting the HMM model with N states, where N = 1 to 5. We checked how the model converged for each value of N by running fitting from 10 random starting states, and chose the N that converged best as the optimum number of states for fitting. Once we choose the best number of states, HMM fitting is done on the full trajectory data, yielding a hidden state sequence for each corresponding trajectory. We used the difference in FLC values between extreme hidden states as a measure of the HMM’s ability to discriminate. Thus, we made a dataset corresponding to different values of 𝑚𝑖𝑛_𝑠𝑎𝑚 𝑝𝑙𝑒𝑠 and 𝜀, and their corresponding best number of HMM states and FLC state difference. We repeated the scheme for all four replicas of each system at different temperatures.

To arrive at an optimized FLC parameter set, we filtered this larger dataset using the following steps: (1) For a given lipid system, select only those pairs with FLC state differences greater than 0.5 (half of the possible range of FLC). This cutoff choice is arbitrary and just enough to identify parameter pairs that distinguish states clearly from each other. (2) From the selected parameter sets, check for the occurrence of a specific pair of (𝑚𝑖𝑛_𝑠𝑎𝑚 𝑝𝑙𝑒𝑠, 𝜀) across the data sets corresponding to all the temperature replicas. Rank the pairs based on most occurrences. This helps to identify parameter pairs that are valid across most temperatures. (3) Choose the pair which gives maximum state difference with most occurances. In case of a tie for the optimum parameter pairs, we chose the pair that has the least 𝜀 and 𝑚𝑖𝑛_𝑠𝑎𝑚 𝑝𝑙𝑒𝑠 to reduce the neighbor search overhead. Now, instead of a temperature-specific 𝑚𝑖𝑛_𝑠𝑎𝑚 𝑝𝑙𝑒𝑠 and 𝜀 for each species in a given ternary mixture, we have a unique pair for each lipid species in a system. We refer to the coordinate using these parameters as FLC_opt_. For the DIPC system, the optimized values (𝑚𝑖𝑛_𝑠𝑎𝑚 𝑝𝑙𝑒𝑠, 𝜀) are (26.5 Å, 24), (31.5 Å, 26), and (32.5 Å, 23) for DPPC, DIPC, and cholesterol respective. For the DAPC system, they are (33.0 Å, 42), (44.5 Å, 56), and (50.0 Å, 41) for DPPC, DAPC, and cholesterol. FLC_opt_ was not used for WE sampling, only as a coordinate on which to project the free energy.

### Weighted Ensemble Simulation

#### Introduction to weighted ensemble simulations

Although the WE method has been reviewed extensively elsewhere(55, 56), we provide a brief overview here. The essence of WE is to generate a large number of short unbiased trajectories, such that they evenly cover conformation space as quantified by a chosen collective variable (CV) or progress coordinate. Specifically, a coordinate is chosen and broken into bins. Short MD trajectories (called walkers) are run, and the location of the final state identified. If a given bin has more than a chosen target number of walkers, some of those walkers are culled and their statistical weight is distributed among the remaining walkers in the bin. If there are fewer than the target number of walkers in a bin (e.g. because more walkers departed than arrived, or because a walker moved into a previously unpopulated bin), the walker is split to produce the target number and its statistical weight evenly distributed among the new walkers.

The WE approach has several major advantages. First, it can be proved that WE converges to the correct free energy distribution in the good-sampling limit (55). Second, its use of a large number of short trajectories is readily parallelizable, making it efficient on high-performance computing resources. Third, it directly produces unbiased estimates of the kinetics that are independent of the chosen coordinate (although some coordinates may converge faster). Fourth, unlike almost every other enhanced sampling method, the MD itself is unbiased; the collective variable is computed only after the MD has finished. This last property was most important to the present application; the coordinates we used are computationally expensive, particularly if a plugin such as PLUMED(87) or COLVARS(52) were used, since both are restricted to a single CPU core and are not GPU-accelerated.

#### Preparing seeding configurations for WE simulation

From each 8 𝜇s MD simulation of a given composition, the last ten frames spaced by 100 ns were collected. These structures contained mixed and unmixed states, which we used as starting structures to begin the weighted ensemble simulations; starting the WE with structures scattered across the collective variable range reduces the time needed to generate well-equilibrated free energy curves.

For the DPPC-DAPC-CHOL and DPPC-DIPC-CHOL systems, the set of mixed configurations for a particular replica came from the respective 423 K and 450 K simulation frames, since both systems are only well-mixed at high temperatures. The set of separated configurations for a replica comes respectively from the 298K and 323K simulations. For the DPPC-POPC-CHOL system, starting structures were taken from the 298K and 450K trajectories; although the system never phase separates, using these two temperatures gives structures that span the whole range of the collective variables.

#### Running WE simulations

We ran weighted ensemble equilibrium simulations using version 1.0 of the WESTPA package(56), closely following the previously established protocol(88). The collective variable was divided into 30 dynamic bins using the minimal adaptive binning scheme (MAB)(89). For each replica, a target number of 4 short simulations, or “walkers” per bin, were started in parallel from the mixed and the separated configurations prepared earlier. After every resampling interval of 1 ns, the collective variable was evaluated to initiate the merging and splitting of walkers to maintain the target number of walkers per bin. One cycle of MD and resampling is referred to as a single WE iteration. We conducted 500 WE iterations for each replica. We used GROMACS 2020.3 to propagate the dynamics, using the same parameters described earlier for the standard MD simulations. The WE Equilibrium Dynamics (WEED) reweighting protocol(90, 91), implemented in WESTPA 1.0, was used to accelerate the convergence of WE walkers toward equilibrium. The reweighting is done every 10 WE iterations. Four independent WE replicas, started with a different subset of initial structures, were simulated for each temperature and lipid composition. All WE simulations were run using the Intel Xeon E5-2695 and Tesla K20Xm GPUs in the BlueHive supercomputing cluster of the Center for Integrated Research and Computing at the University of Rochester.

#### Analysis of WE simulations

The probability distributions for the CVs for each replica, as a function of WE iterations, were constructed using 𝑤_𝑝𝑑𝑖𝑠𝑡 and 𝑝𝑙𝑜𝑡 ℎ𝑖𝑠𝑡 tools in WESTPA. Using this distribution, we monitored the evolution of each WE replica simulation and the convergence. We used the 𝑤_𝑚𝑢𝑙𝑡𝑖_𝑤𝑒𝑠𝑡 tool in WESTPA to combine data from four the WE replicas of a system at a given temperature. We then constructed the respective free energy surface from the combined probability distribution of a system.

Unless otherwise noted, equilibrium curves were always calculated using the last 10 iterations of each WE run, because combining results pre- and post-reweighting is not statistically correct. To check the populations in different states and the determine flux between states the 𝑤_𝑖 𝑝𝑎 tool was used. When performing this calculation for the DPPC-DIPC-CHOL system at 323K, we defined the states based on visual inspection of the corresponding free energy curves. We defined the mixed and separated states as CEI = [0.0, 3.9] and [4.4, 6.0], SI_noCHOL_ = [0.0, 1.3] and [1.4, 2.0], and FLC = [0.0, 0.575] and [0.65, 1.0] respectively. The choice is arbitrary but reasonable enough to give us a picture of what is happening.

Input files, initial structures, and scripts required to run and analyze all simulations are available on GitHub at https://github.com/Poruthoor/Phase_Separation_Article/tree/main/FLOPSS

## RESULTS

Consistent with previous studies from which they are adapted, the standard CG MD simulations of DPPC-(DA/DI)PC-CHOL systems phase separates into L_o_ and L_d_ regions. The L_o_ region is enriched in the saturated lipid, DPPC, and cholesterol, while the L_d_ region contained mostly the unsaturated lipids, DAPC or DIPC, depending on the system. As expected, the DPPC-POPC-CHOL system did not form distinct phases based on visual inspection. This section compares how different variables track phase separation propensity in lipid bilayers using standard CG MD simulations. We then compare the convergence of WE simulation to the choice of collective variable. Finally, we present the free energy landscapes of lipid bilayer systems obtained using WE simulations and discuss reusing the data generated to form other intuitions and applications.

To evaluate how the collective variables track phase separation, we traced the time evolution of each variable for each system using standard MD starting from well-mixed bilayers. Figure 3 illustrates the temporal evolution of FLC, CEI, and SI_noCHOL_ for all systems at 298K, 323K, 423K, and 450K. For the two phase separating systems, the variables capture a single transition between a mixed state and a separated state. Interestingly, for the relatively slow-separating DPPC-DIPC-CHOL system, CEI and SI_noCHOL_ appear to indicate a slower transition than FLC, indicating that while these variables are correlated, they do not measure precisely the same phenomenon. For the DPPC-POPC-CHOL system, the variables capture a single state corresponding to a mixed system and no transition. Nevertheless, it is worth noting that FLC, CEI, and SI_noCHOL_ capture the effect of temperature in all systems, including for POPC, the negative control, that does not phase separate.

**Figure 3:**
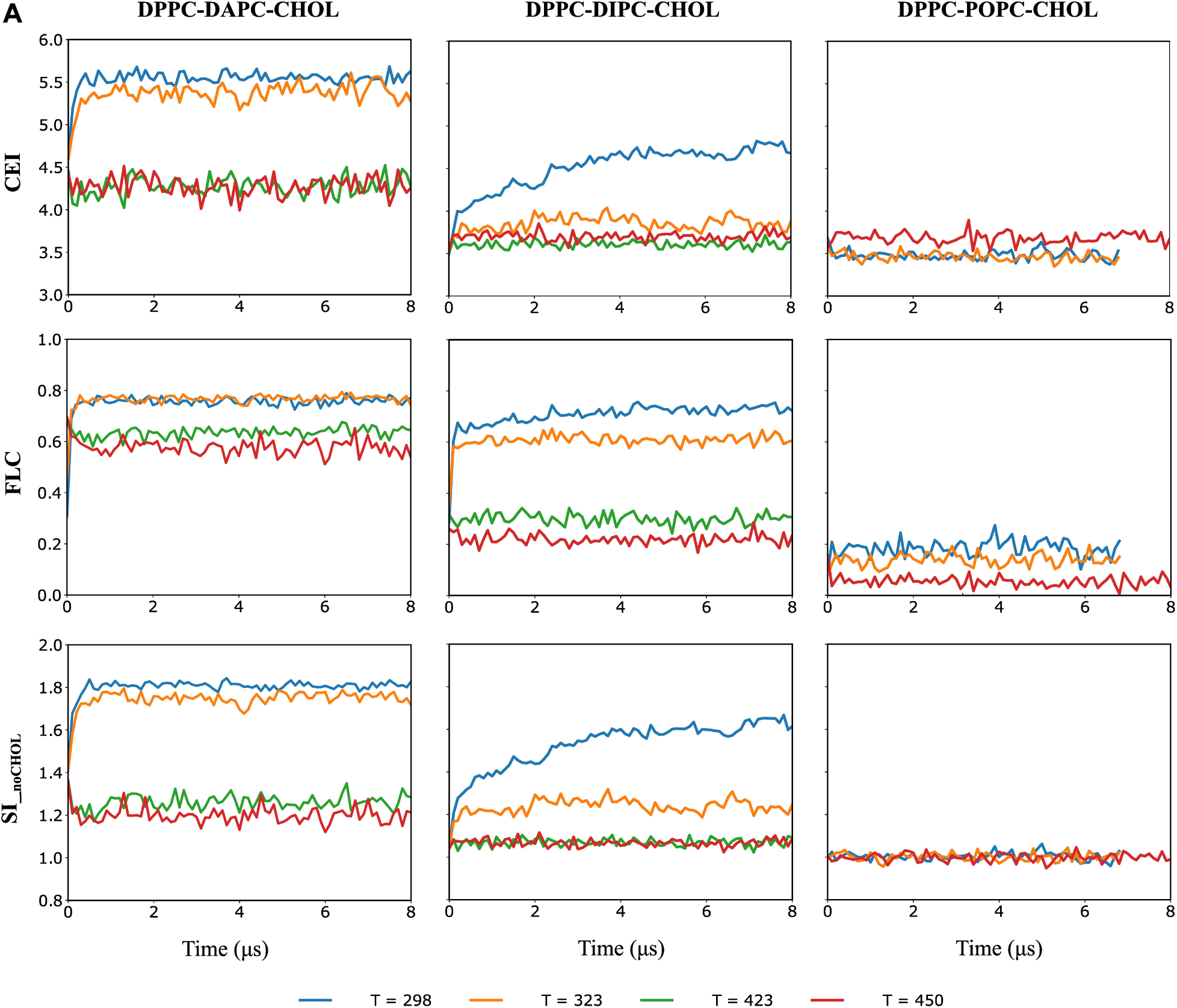
A. Time series for three collective variables tracking membrane separation. Each line represents one conventional MD simulation started from a well-mixed state. Each column contains trajectories from a single lipid composition, while each row shows a different variable; 8𝜇s trajectories were used for all rows. The lines are colored to represent different simulation temperatures.

These variables even capture the subtle differences in phase-separating propensity between systems. For example, at 298K, the plateaued region of the FLC, CEI, and SI_noCHOL_ curves are higher for the DPPC-DAPC-CHOL system than the DPPC-DIPC-CHOL system. This is expected as (a) the lipid chain mismatch between saturated and unsaturated lipid species and (b) the number of double bonds in unsaturated lipid species is high in the DPPC-DAPC-CHOL compared to DPPC-DIPC-CHOL system. Both these factors have previously been shown to influence lipid phase separation kinetics and domain stability(35, 92). Thus we have a set of low-dimensional variables that can (a) represent the global dynamics of the lipid bilayer system, (b) distinguish and track the transition between mixed and separated states, and (c) capture temperature effects, based on the composition of the system.

### Choice of collective variables for WE simulations

Since CEI, SI_noCHOL_, and FLC all track bilayer lipid separation similarly, we decided to test them as progress coordinates to drive the WE simulation. Much to our surprise, the 3 coordinates perform very differently. Figure 4A shows free energy curves computed for each coordinate using the DPPC-DIPC-CHOL system at 323K; each curve on a given plot was computed from a different block of 10 WE iterations for the same simulation. The free energy curves generated using CEI and SI_noCHOL_ are noisy and not self-consistent, indicating that at best there is poor statistical convergence despite the relatively lengthy 500-iteration WE runs. The SI_noCHOL_ runs are still more troubling, in that they suggest that the well-mixed state is favored over the separated one, which contradicts the results from the conventional MD simulations. By contrast, the free energy curves generated using FLC as the collective variable are quite self-consistent, with the phase-separated state slightly favored. Interestingly, if we monitor the evolution of the free energy curves as a function of WE iterations, shown in Fig 4B, the simulations appear to have converged in the sense that they are no longer systematically changing.

**Figure 4:**
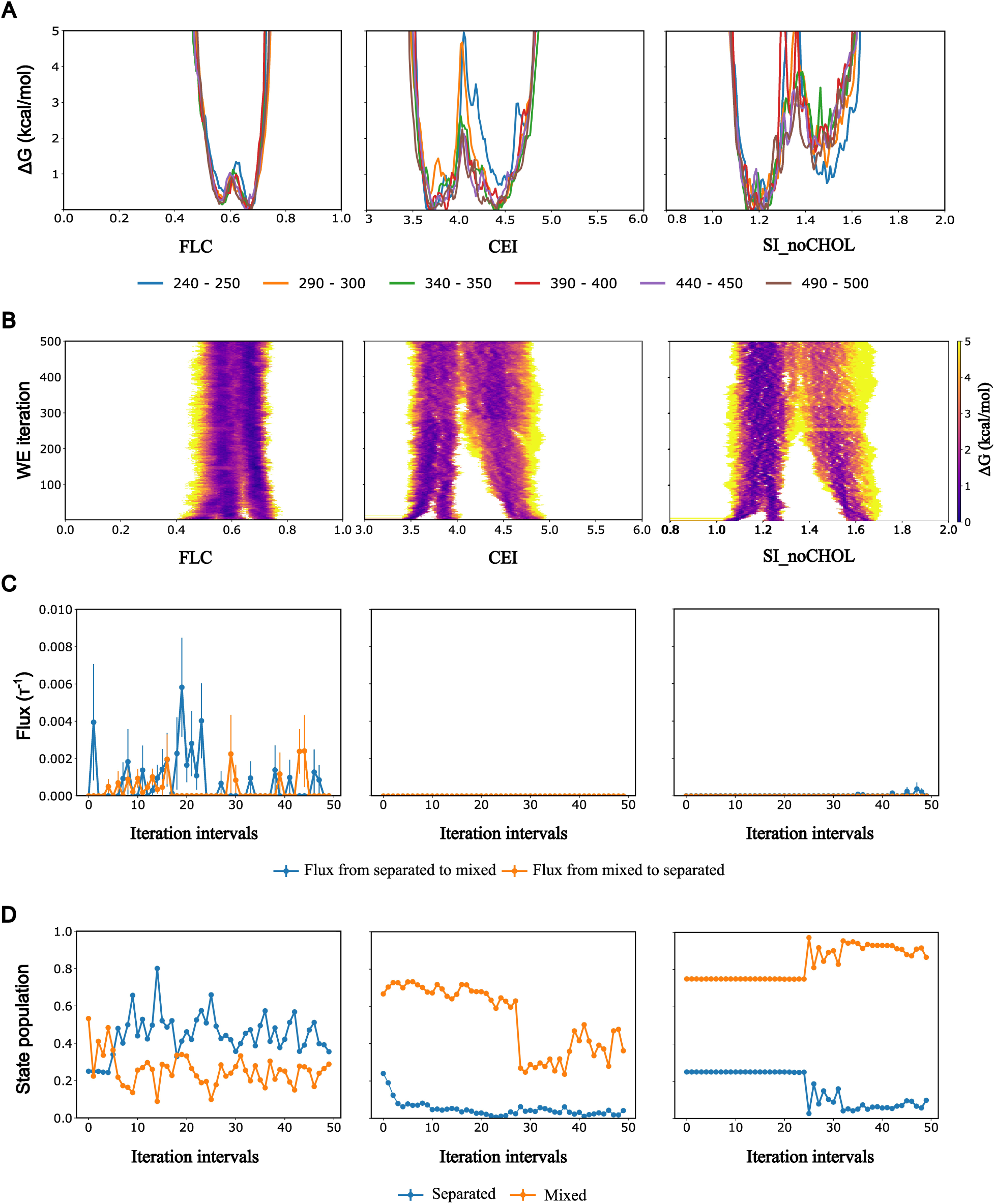
For the given DPPC-DIPC-CHOL replica: A. Free energy curves as a function of WE iteration blocks used to generate them. B. Each plot gives the evolution of configurational distribution for a given collective variable. C. Flux profile between states. The peaks represent crossing of walkers. D. Corresponding state population. Here, each column represent the collective variable that was used to drive WE simulation (FLC, CEI, and SI_noCHOL_).

These conflicting results raise the following questions: Why do variables that make sense while tracking the system in standard CG MD simulation yield inconsistent or incorrect free energy landscapes when used as collective variables in a WE simulation? One clue to diagnose the problem lies in our starting states. In contrast to most WE calculations, which begin the simulation with all walkers in the “first” bin, we started with diverse states from across the whole range of the collective variable. While the free energy curves evolve as the WE proceeds, for the poorly performing coordinates the relative probability of the 2 states remains almost entirely unchanged.

Figure 4C confirms this hypothesis by tracking the probability flux between the two states for each coordinate; the results confirm that when CEI or SI_noCHOL_ are used as progress coordinates, there is essentially no flux across the barrier separating the separated and mixed states; the flux is literally 0 for CEI, while SI_noCHOL_ shows very small fluxes during the last 100 iterations. By contrast, WE simulations using FLC as the coordinate consistently show flux between the two states that is at least several orders of magnitude larger.

Figure 4D shows this information another way, tracking the relative population of the mixed and separated states over the course of the WE runs; for clarity purposes the states are defined to exclude the “middle” of the coordinate (see the methods section for details), so the state populations do not necessarily add to 1. It is interesting to note that for CEI, we see a significant drop in the population of the mixed state during iterations 290-300, without a concomitant rise in the other state. Since our state definition we chose does not span the entire collective variable space leaving a room for the barrier that separates two states, this indicates that some walkers transitioned out of the mixed state, but remained trapped in the middle of the collective variable and never transitioned to separated state.

For reasons that are not entirely clear, one coordinate (FLC) is far more efficient at sampling transitions than the other two. CEI and SI_noCHOL_ do not generate crossing events, and as a result largely recapitulate the probability distribution used to seed the calculation. The combined results from standard CG MD and WE simulations suggest that CEI and SI_noCHOL_ are suitable proxy labels for phase separation but perform poorly as collective variables for sampling. For this reason, we only continued testing the FLC-based WE calculations for other systems and temperatures.

Figure 5 shows the free energy profiles of the DPPC-DAPC-CHOL, DPPC-DIPC-CHOL, and DPPC-POPC-CHOL lipid bilayer systems. Each curve is generated by averaging four WE replica simulations, as described in the Methods section, using the last 10 iterations (491–500) of each replica. The variance between the individual replica free energy curves is generally quite small, approximately 0.5 kcal/mol, shown in Supplementary Figure S7. The DPPC-DAPC-CHOL system has a double well behavior at 323K and 353K. However, both basins correspond to high FLC, such that the lower FLC basin is still extremely non-ideally mixed. Based on visual inspection, it seems the primary difference is the degree to which lipids have coalesced into domains. Indeed, even at 423 K, we see that this system prefers to be in configurations where more than 60% of lipids are in clusters. For the DPPC-DIPC-CHOL system, the double-well free energy curves gradually shift to the left (fewer lipids in clusters) and change to narrower single-well curves as the temperature increases. For the DPPC-POPC-CHOL system, the free energy curves at 298K and 423K have a single basin and correspond to a low fraction of lipids preferring to be in any clusters. In general, FLC-based free energy landscapes capture the role of lipid species constituting the bilayer system in its phase separation. Moreover, the effect of temperature in decreasing the propensity of a lipid bilayer to separate is evident in all systems. Although the expectation of clustering-based FLC is to track the formation of domains in a lipid bilayer, the fact that it also neatly captures the temperature effects, even for the negative control POPC system, suggests the robustness of FLC as the collective variable.

### Reconstructing free energy landscapes on alternative coordinates

One outcome of WE simulation is the ensemble of diverse structures generated by the trajectories. By construction, WE resampling ensures that the weights associated with these walkers are known and unbiased. Thus, we can reuse these weights to examine other variables of choice post-simulation without any need to reweight; this is advantageous, because reweighting nearly always introduces significant noise due to heavy weights applied to the lowest Δ𝑈 structures. Assuming the initial configurational distribution relaxed into an equilibrium distribution during the WE simulations, we can reconstruct analogous free energy landscapes with collective variable candidates that underperformed for sampling but are otherwise more intuitive to understand. Fig 6 shows reconstructed free energy landscapes using 3 coordinates, each constructed using the structures and weights from last 10 iterations of each FLC-generated replica; Panel A shows the results for SI_noCHOL_, Panel B shows the results for CEI, and Panel C shows the results projected onto FLC_opt_. The results are qualitatively similar on each coordinate, though unsurprisingly there are quantitative differences. A few features are worth noting: First, all 3 coordinates show more distinct structure than the original FLC-based free energy curves. The DIPC system at low temperature in particular produces free energy curves with significant fine structure, although it is unclear whether that structure is merely the result of statistical noise. Second, FLC_opt_ produces far more spread out curves than the original FLC, which is consistent with the idea that FLC_opt_ is a more sensitive coordinate. With the exception of the systems at 423K, all of the free energy curves have at least two well-defined wells.

**Figure 6:**
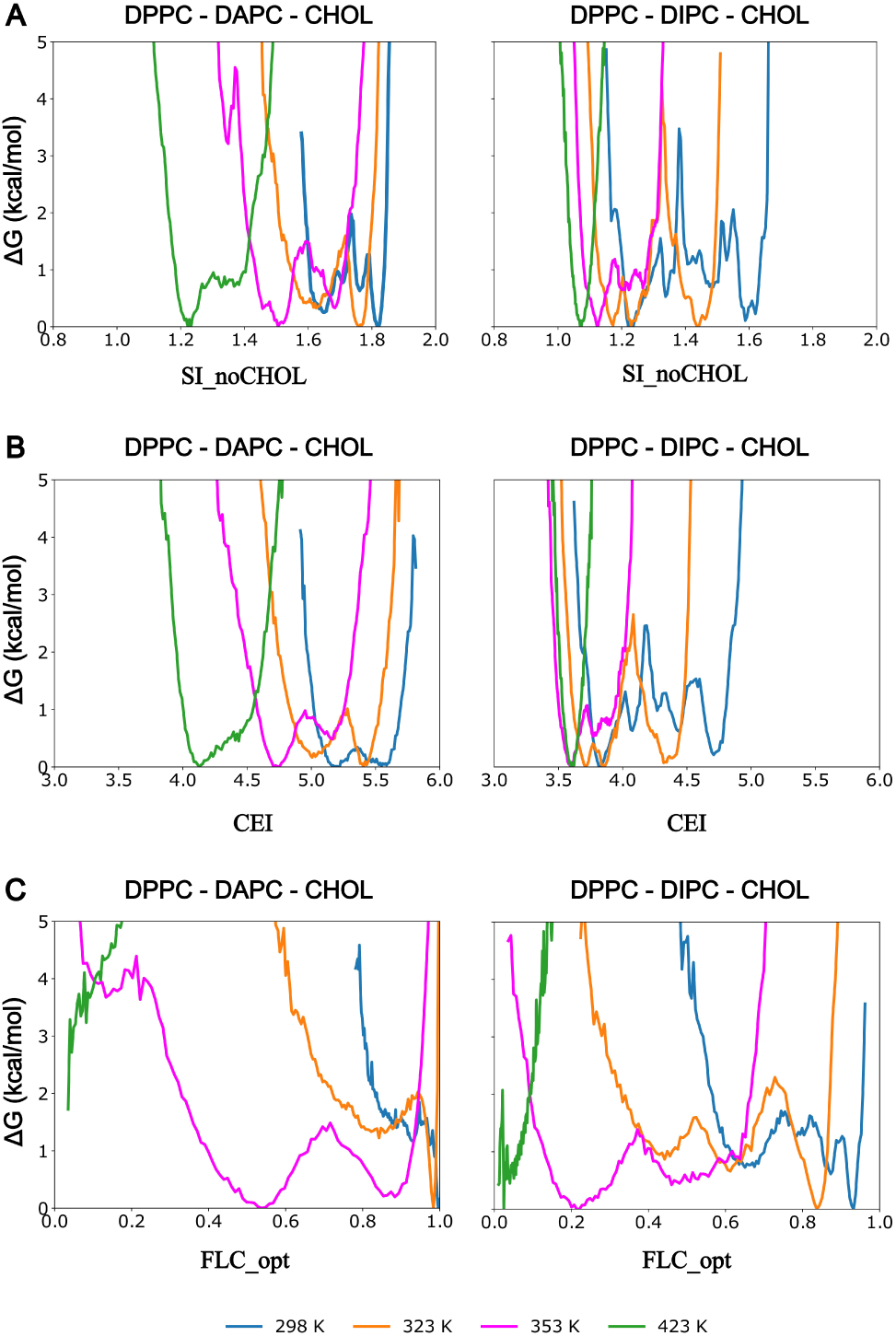
A. Free energy landscapes reconstructed using the ensemble of configurations generated during last 10 iterations of FLC-driven WE simulation. The free energy landscape is reconstructed with respect to SI_noCHOL_ for DPPC-DAPC-CHOL and DPPC-DIPC-CHOL system respectively. B. The free energy landscape is reconstructed with respect to CEI for DPPC-DAPC-CHOL and DPPC-DIPC-CHOL system respectively. C. The free energy landscape is reconstructed with respect to FLC_opt_ for DPPC-DAPC-CHOL and DPPC-DIPC-CHOL system respectively. In all panels, the lines are colored to represent different simulation temperatures.

### Computing the free energy of phase separation

If we define a cutoff FLC value that separates the mixed and phase-separated states, we can compute the free energy change for phase separation as

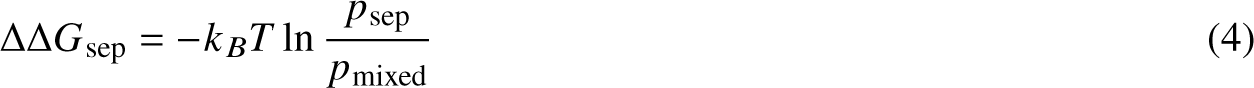

where the probabilities are extracted by summing the weights from the weighted ensemble simulations.

One challenge, however, is that the locations of the basins shift with temperature, as seen in Fig 5A, and at higher temperatures, the notion of two states — mixed and separated — breaks down as the double-well behavior of free energy curves turns into a single well. One trivial solution would be to define a rigid FLC cutoff for a system. However, a caveat for this approach, as seen in Fig 5 A, is that the free energy basin for a specific state of the DPPC-DIPC-CHOL system is different for the respective state basin for the DPPC-DAPC-CHOL system. For this reason, devising a rigorous systematic way to define the separator between the states will require more investigation. As a first attempt, we use visual inspection to define FLC cutoffs of 0.525 and 0.625 for the DPPC-DIPC-CHOL and DPPC-DAPC-CHOL systems respectively; below those values, the systems are relatively well-mixed. Figure 7 shows the ΔΔG curve for the DPPC-DAPC-CHOL and DPPC-DIPC-CHOL systems as a function of temperature for respective FLC cutoff. In principle, one can identify a melting temperature 𝑇_𝑚_ as the temperature at which ΔΔ𝐺_sep_ = 0; however, the value derived in this manner is sensitive to the choice of FLC cutoff, as shown in Supplementary Figure S8.

**Figure 5:**
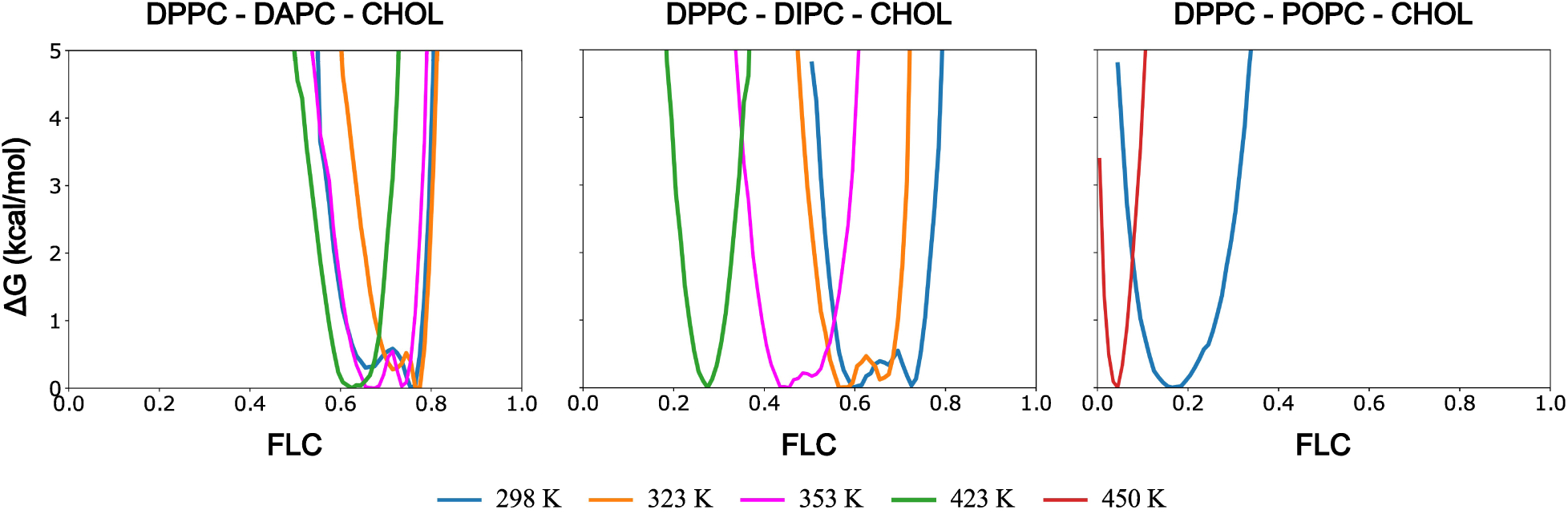
Free energy curves based on FLC-driven WE simulation for three systems. Each line represents a free energy curve, computed as the average of 4 replicate WE runs. Each column contains free energy curves from a single lipid composition. The lines are colored to represent different simulation temperatures.

**Figure 7:**
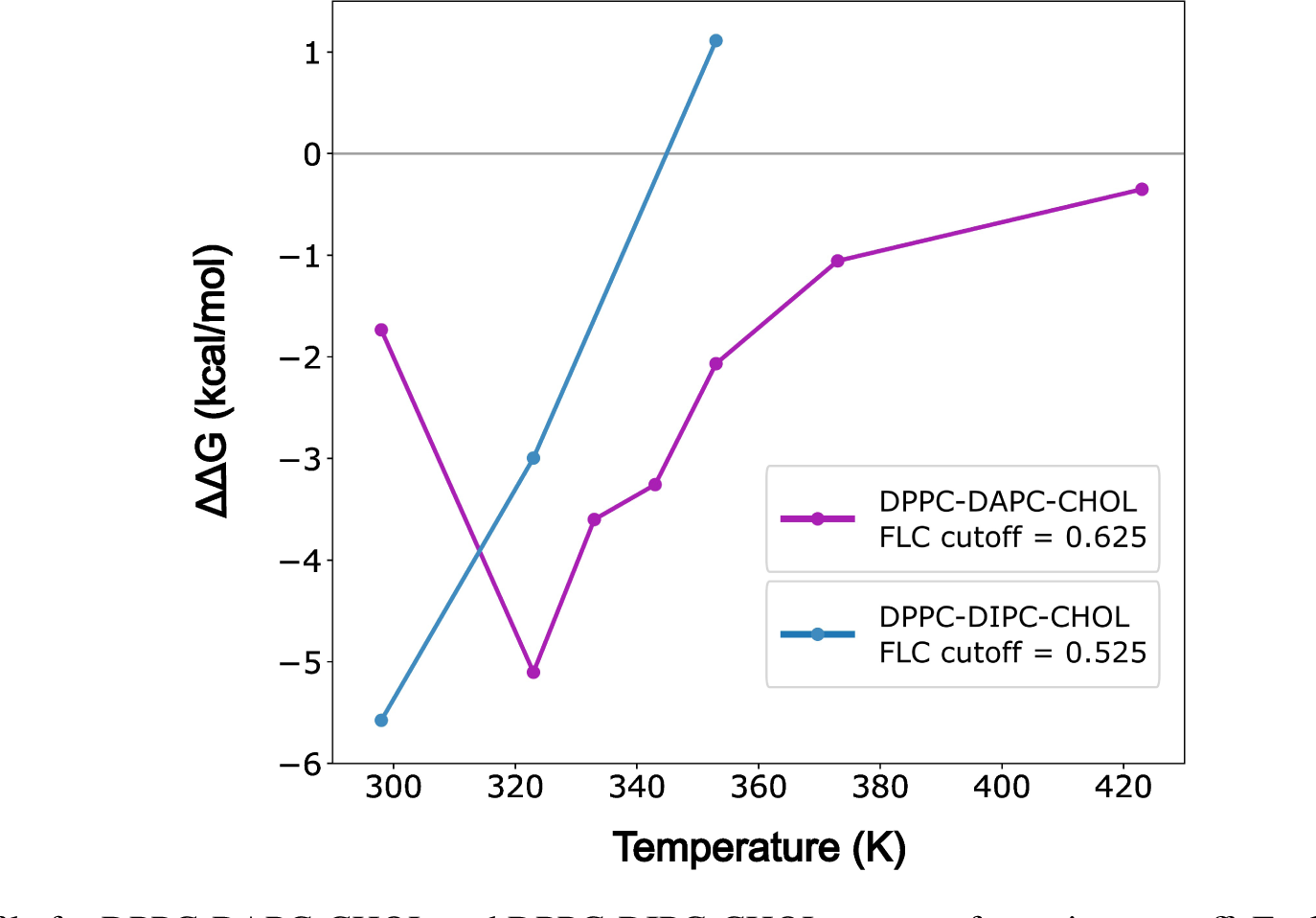
ΔΔG profile for DPPC-DAPC-CHOL and DPPC-DIPC-CHOL systems for a given cutoff. Each curve represents ΔΔG for the system to transition from mixed to the separated state. Negative values indicate phase coexistence is favored. Figure S8 shows the sensitivity of the computed ΔΔG to the choice of cutoff.

An arguably more satisfying approach is presented in Figure 8. Here, we take advantage of the presence of more structure in the free energy as a function of FLC_opt_ compared to the original FLC. The presence of well-separated basins in the free energy curves lends itself to identifying the local maxima as boundaries between states; with this distinction drawn, we can compute the free energy difference using Equation 4. This breaks down for the 423K systems. Because these curves have only a single well, we are forced to identify a cutoff value; the plotted values should be considered a floor for the “true” value.

**Figure 8:**
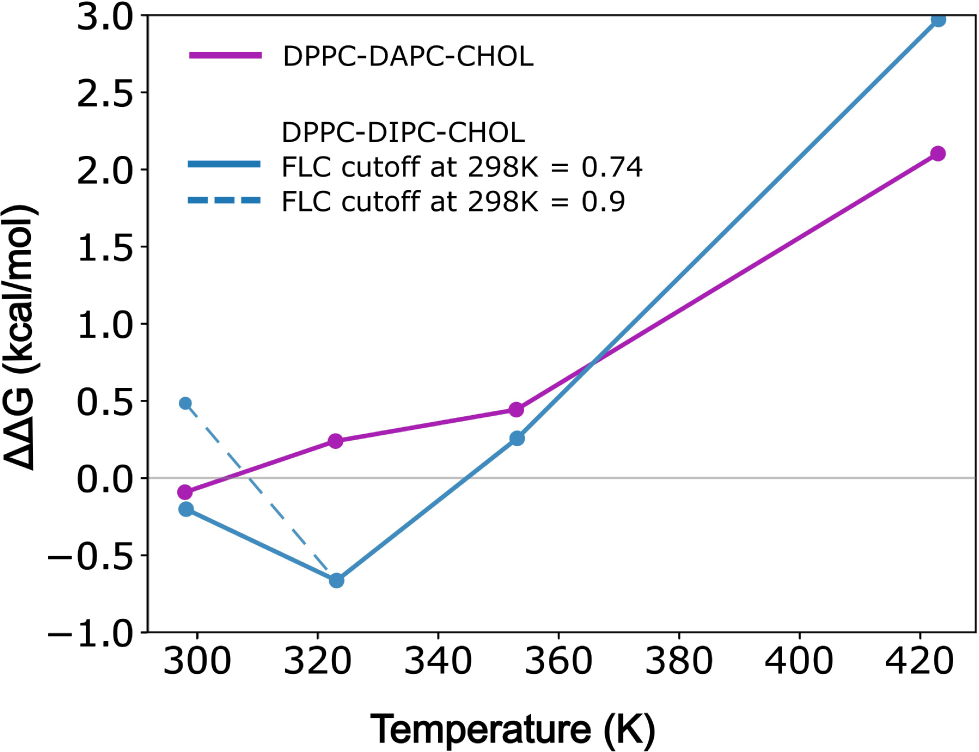
ΔΔG profile for DPPC-DAPC-CHOL and DPPC-DIPC-CHOL systems constructed by separating FLC_opt_ into 2 states at the first local free energy maximum from the right. For the DIPC system at 298K, there are more than 2 basins, and as a result there are 2 plausible places to separate the states, at FLC_opt_ values of 0.9 and 0.74 (see Figure 6); the latter value is marked with a dashed line. Negative values indicate phase coexistence is favored.

## DISCUSSION

### Relative effectiveness of collective variables for sampling

It is well-established that the choice of collective variable is crucial for the success of enhanced sampling protocols(51, 76, 77). However, determining in advance which coordinates are best suited to efficient sampling is far from obvious. Several groups, most notably that of Tiwary (93–96), have developed machine learning methods to identify effective coordinates empirically based on their kinetic properties. A more theoretical approach was developed by Wu et al (97), based on identifying the energy flow coupled to motion along a coordinate. However, the former methods do not explain the reason why a particular coordinate is effective, while the latter is too computationally intensive to apply to systems like those treated here; the presence of a large number of equivalent molecules is a significant challenge to the formalism as currently presented.

Thus, we are left with less-satisfying intuitive explanations for the wide variation in performance of the collective variables tested here. It is reasonable to assume that the most efficient coordinate for sampling is one that closely approximates a “true” order parameter for the system, and thus that the efficacy of FLC as a sampling coordinate says something about local clustering as a mechanism of phase separation. In particular, FLC is sensitive to somewhat longer lengthscale motions than CEI or SI, both of which are driven entirely by nearest-neighbor mixing. This is also borne out by the improved ability of FLC_opt_ to discriminate between states; the optimized cutoff distances are roughly 2-3x longer. Based on this, we would predict that FLC_opt_ would perform as well or better than FLC, though this will require more testing.

#### **Interpreting** ΔΔ𝐺**_sep_**

It is tempting to think of ΔΔ𝐺_sep_ the way we do with simple two-state double-well systems, where the relative populations of the two states change with temperature but the nature of those states is unchanged. Consideration of Figures 5 and 6 show that is clearly not the case here; regardless of the collective variable used to project the free energy, the location of the minimum or minima varies strongly with temperature. This is even visible in the non-separating POPC system, where the free energy curve shifts toward lower FLC at higher temperatures (Figure 5) This is not surprising and has a direct physical interpretation: above the 𝑇_𝑚_, there is a single phase, but the lipid do not mix ideally; this is most visible in the DAPC system, where most DPPC lipids are clustered even at 423K. Even below the melting temperature, the structure and composition of the phases changes with temperature.

As a result, the computed ΔΔ𝐺_sep_ values (Figures 7 and 8) are not nearly as large as one might expect. For example, the separated state for the DIPC system at 298K is about -5 kcal/mol when estimated by choosing a conventional FLC cutoff that “looks” reasonable. However, if we use FLC_opt_ and choose to split the states at the first free energy barrier from the right (high FLC_opt_), we get a much smaller difference of -0.74 kcal/mol. The difference is that the former approach attempts to compare the free energies of a relatively well-mixed state to the separated one, while the latter compares the two stable states of the system: one phase-separated, and the other a single phase with very non-ideal mixing. Since at low temperature good mixing is very unfavorable, the latter approach produces a far smaller ΔΔ𝐺_sep_. That said, it is also possible that this relatively moderate favorability is an artifact of the size of the system simulated; in the future, it would be worthwhile to investigate the dependence of ΔΔ𝐺_sep_ on system size in a more systematic way.

This explanation also rationalizes the most surprising result, that the DIPC system has more favorable ΔΔ𝐺_sep_ than the DAPC system at all temperatures except 423K, even though we would have thought of DAPC as being the more strongly separating system. However, once we consider the fact that the DAPC is far more non-ideal when existing as a single phase than the DIPC system, the result makes more sense: the non-ideality means that the DAPC system is more able to sequester the DPPC lipids without the entropic penalty of forming larger domains. Put another way, the change in local composition surrounding a given DPPC is smaller upon phase separation for the DAPC system than for DIPC. That said, it is also possible that system-size dependence is playing a role here as well; for example, if DAPC domains naturally existed on longer lengthscales than DIPC, that would explain the results.

### FLOPSS pipeline for computing phase separation thermodynamics

Here we propose a proof-of-concept pipeline to construct **F**ree energy **L**andscape **O**f **P**hase **S**eparating **S**ystems (**FLOPSS**) by realizing multiple transition events using WE strategy. Even though our systems of interest are lipid bilayers, using appropriate model resolution and collective variables can potentially generalize this pipeline to any system that phase separates. The modular pipeline is outlined in Figure 9, with different sublayers that constitute each module.

**Figure 9:**
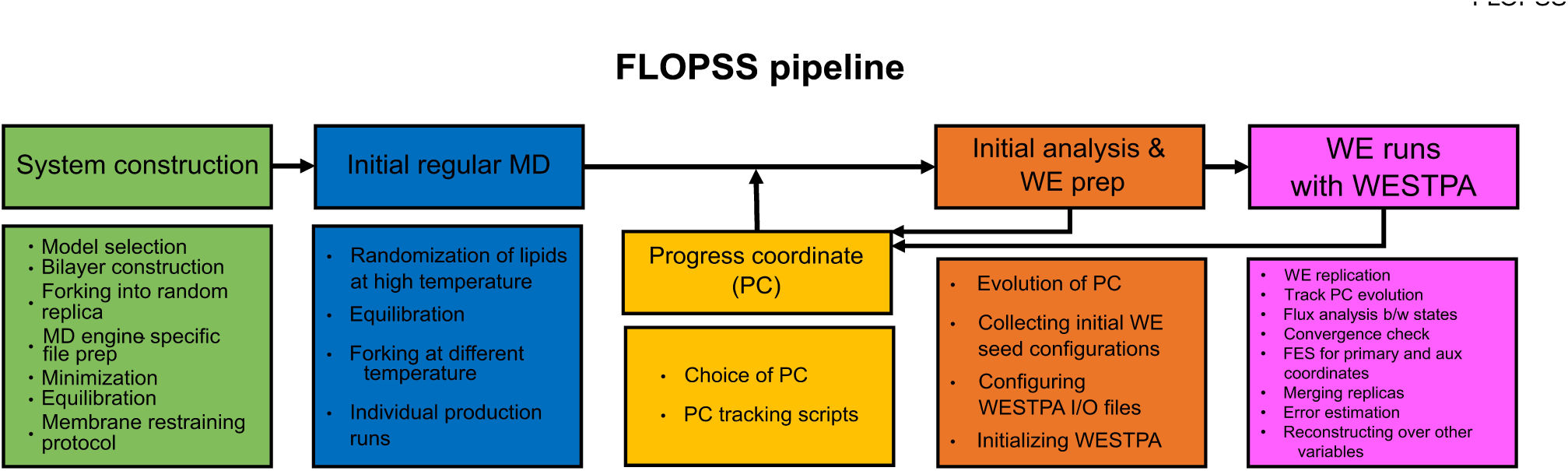
Schematic representation of the proposed FLOPSS pipeline. The pipeline is modular. Each module has layers that can be translated to any phase-separating system.

The free energy landscapes generated with FLOPSS are consistent with behaviors from the literature, opening up applications to other systems, including asymmetric bilayers, bilayers including peptides or proteins, or all-atom models. We also plan to investigate the effects of finite-sized systems on the phase separation thermodynamics (98). One additional advantage of basing the method on WE is that it can also be used to study the kinetics of phase separation as well as the thermodynamics. One other major area of research is determining whether other collective variables might perform better and allow for less ambiguous procedures for determining ΔΔ𝐺_sep_. Finally, many of the details of the pipeline could be further optimized, especially those associated with the WE runs themselves.

## CONCLUSION

In this work, we have proposed a simple yet efficient collective variable that simultaneously tracks phase separation and drives WE simulation, ensuring sufficient state crossing with reasonable convergence of configurational distribution. We give yet another example of why collective variable choice is crucial for the success of enhanced sampling protocols, and that there can be considerable gaps in performance between otherwise reasonable variables. Thus, a more thorough and systematic analysis of the sensitivity of WE simulation on the choice of collective variable is needed and is unfortunately beyond the scope of this work.

In summary, we have developed and validated a new framework that can directly compute the thermodynamics associated with lipid phase separation from simulation. We have also demonstrated the potential reuse of a reasonably well-converged WE simulation driven by a good collective variable to explore other variables, which otherwise constitute a poor choice for driving WE simulation. Thus, we can increase the effectiveness of WE simulation without compromising on computational cost. Moreover, we also showcase the potential of FLOPSS to construct a ΔΔG profile of the system under study to investigate the melting properties. We want to highlight that by generalizing the collective variable FLC to track clustering in 3D space, FLOPSS can in principle be extended to other instances of biological phase separation.

## AUTHOR CONTRIBUTIONS

AJP and AG designed the project. AJP and AS performed the research. AJP and AG wrote manuscript.

## ACKNOWLEDGEMENTS

This work is dedicated to Dr. Klaus Gawrisch, who has inspired much of our interests in lipid bilayers. The authors thank the Center for Integrated Research Computing (CIRC) at the University of Rochester for providing computational resources and technical support. The authors also thank Prof. Lillian Chong and Prof. Daniel Zuckerman and the WESTPA Data and Dev club for insights. The authors thank Anthony Bogetti, Jeremy Leung, John Russo, Dr. Sreyoshi Sur, and Dr. Tod D. Romo for their support. This work was supported by grant R21GM138970 (to AG) from the National Institutes of Health.

