## Supplementary material for "Understanding the Free Energy Landscape of Phase Separation in Lipid Bilayers using Molecular Dynamics": 9 supplemental figures and text

August 25, 2023

### 1 Simulation details[†]

| | DPPC-DAPC-CHOL | DPPC-DLiPC-CHOL | DPPC-POPC-CHOL |
|---|---|---|---|
| Bilayer composition | 600:360:240 (0.5:0.3:0.2) | 828:540:576 (0.42:0.28:0.3) | 480:480:240 (0.4:0.4:0.2) |
| Water[‖] | 9000 PW Beads | 14580 PW Beads | 9000 PW Beads |
| Temperatures for std. MD production runs | 298K, 323K, 333K, 333K, 343K, 353K, 373K, 423K, 450K | 298K, 323K 353K, 423K 450K | 298K, 323K, 450K |
| Temperatures for WE MD production runs | 298K, 323K, 333K, 333K, 343K, 353K, 373K, 423K | 298K, 323K 353K, 423K | 298K, 450K |

| | |
|---|---|
| Temperature | Initial NPT equilibration @ 400 K, with Berendsen for 100 ns. Production run at with v-rescale, $\tau$ = 1.0 ps |
| Time step | 20 fs |
| vdW | Potential-shift-verlet. Cutoff = 1.1 nm. |
| Electrostatics | Reaction field. Cutoff = 1.1 nm Rel.dielectric constant = 2.5 (Since we are using polarizable water) |
| Pressure | Parrinello-Rahman @ 1 bar. Semi isotropic. $\tau$ = 12 ps. Compressibility = 3 $\times 10^{-4}$ |

Table 1: Simulation details. [†] The system parameters were obtained from the .mdp file for production stage which is given by CHARMM GUI[1] after building the system. These parameters agree with the MARTINI recommended parameters[2] for a CG lipid system that uses polarizable water[3]. [‖] PW : Polarizable Water.

#### 2 Membrane restraint protocol

Membrane undulation is a known behavior in large lipid bilayer systems; when undulations are undesirable (as in our case) membrane restraint protocols have been used to prevent them[4, 5, 6]. There are two common protocols that have been used previously: (a) position restraining a certain bead of specific lipid or (b) applying a flat bottom restraint on the membrane. For detailed explanation of how this is implemented, refer to the GROMACS manual[7].

We chose flat-bottom restraints, where the lipids can move in the xy slab of predefined z thickness without any perturbation, because it is a smaller perturbation. Thus, starting from the last 100 ns of NPT equilibration of lipid bilayer to the whole production run, we introduced a flat-bottom restraint to the bilayer.

There are two parameters to choose: (a) How thick should be the planar slab where bilayer is free to move, and (b) What should be the force constant to be used for the flat bottom potential In order to decide on the former, we found the mass density distribution of NC3, PO4 and GL1 beads along z direction during the CHARMM-GUI relaxation protocol. Based on the distributions at equilibration step, for the DPPC-DIPC-CHOL, we chose 2.3125 nm as the flat bottom well thickness, with respect to PO4 bead of DPPC and DIPC. For the DPPC-DAPC-CHOL, we chose 2.5625 nm as the flat bottom well radius, with respect to PO4 bead of DPPC and DAPC. We chose PO4 bead since the head group choline (NC3) seemed more flexible (as instead of a sharp peaks at each leaflets it has a more broader peaks at each sides of the plots), therefore for the NC3 bead average value has more uncertainity to it. Also, PO4 bead to bead distance for DPPC and D(A/I)PC was different but we went with the idea of one height for both, at the outer edge of the farther one. Also we only applied the flatbottom restraints for the longer DPPC and D(A/I)PC lipids and not for cholesterol. We also tested flat bottom restrain radius = 21 A, 22 A, 23 A, 24 A, 25 A, 26 A, 27 A for DIPC system and checked whether the criteria we chose prevented bilayer undulation while not otherwise stiffening the bilayer. Based on visual inspection, it seems our choice of criteria was effective.

We tried flat bottom restraints with force constant, $k = 0.2, 2, 10, 20, 100, 200, 500, 1000, 2000, 3000$ KJ mol-1nm-2 for DIPC system. We ran 100 ns NPT runs and observed that force constants below 1000 mol-1 nm-2 still allowed undulations. Based on this, as well a review of recent literature [6], we chose $k = 1000$ KJ mol-1 nm-2 for DIPC system.

### 3 Radius cutoff calculations for $\epsilon$

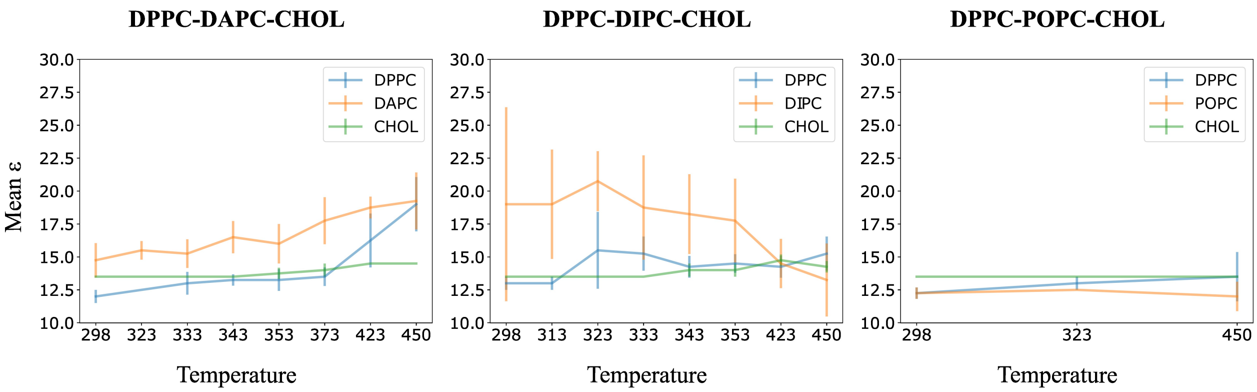


Figure S1: Mean $\epsilon$ calculated using the *xy_rdf* tool in LOOS package at different temperature and for different lipid species in each system under study.

### 4 Auxiliary variables

To assess the quality of DBSCAN clustering, for each lipid species $X_i$ in the system, we calculated the following,

1. Number of $X_i$ clusters in the system under study.
2. Fraction of $X_i$ lipids in $X$ clusters
3. Fraction of $X_i$ lipids in $X$ core lipids.
4. Mean Silhouette Coefficient (MSC) of $X_i$ Clusters, as implemented in scikit-learn.

The Silhouette Coefficient, a method used to evaluate the clustering done by any technique, is defined as

$$\mathrm{s} = \frac{b - a}{max(a, b)} \tag{1}$$

Here, for a $X_i$ lipid in the cluster, $a$ is the mean inter-lipid distance for $X_i$ lipids within a given cluster, and $b$ is the mean minimum distance between clusters. The former assesses the 'cohesion' of a given $X_i$ lipids with other $X_1$ lipids in a cluster, while the latter assesses the 'separation' from the nearest cluster.

The Mean Silhouette Coefficient of $X_i$ Clusters is given by the mean $s$ over all non-outlier $X_i$ lipids. Here, we have omitted the MSC calculations for cases when there are no clusters or just one cluster detected by DBSCAN. MSC is bound between and -1 and 1. A high positive value corresponds to well-segregated dense clusters, while a low negative value implies that lipids are assigned to clusters incorrectly.

All these AVs were tracked along with the primary collective varibale, FLC, driving the WE simulation. Following are the system specific profiles of each AVs:

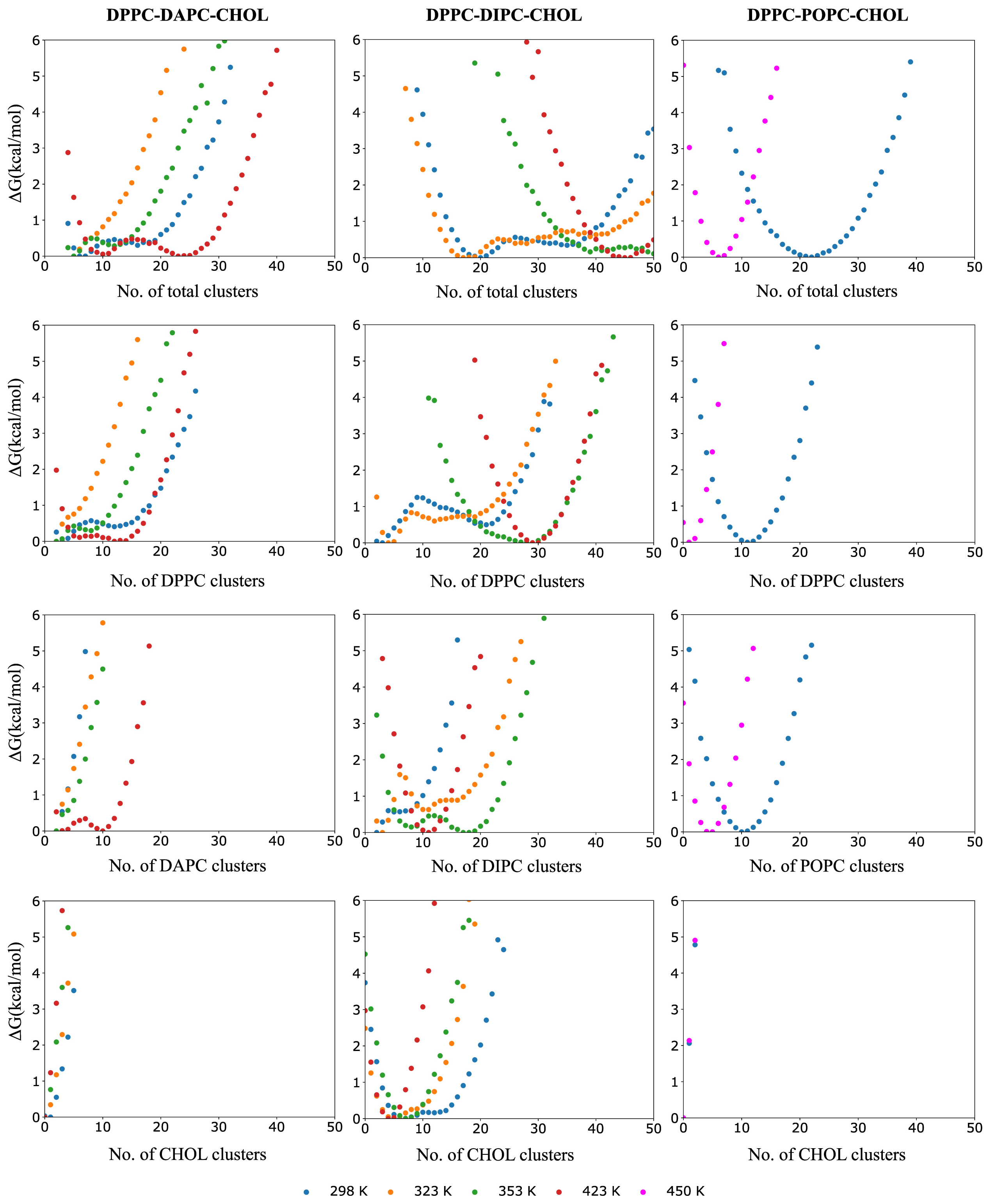


Figure S2: Cluster count profile for each system as a function of temperatures. Each column represents each lipid system under study. The rows represent the total cluster count, saturated lipid cluster count, unsaturated cluster count and cholesterol cluster count within each system respectively

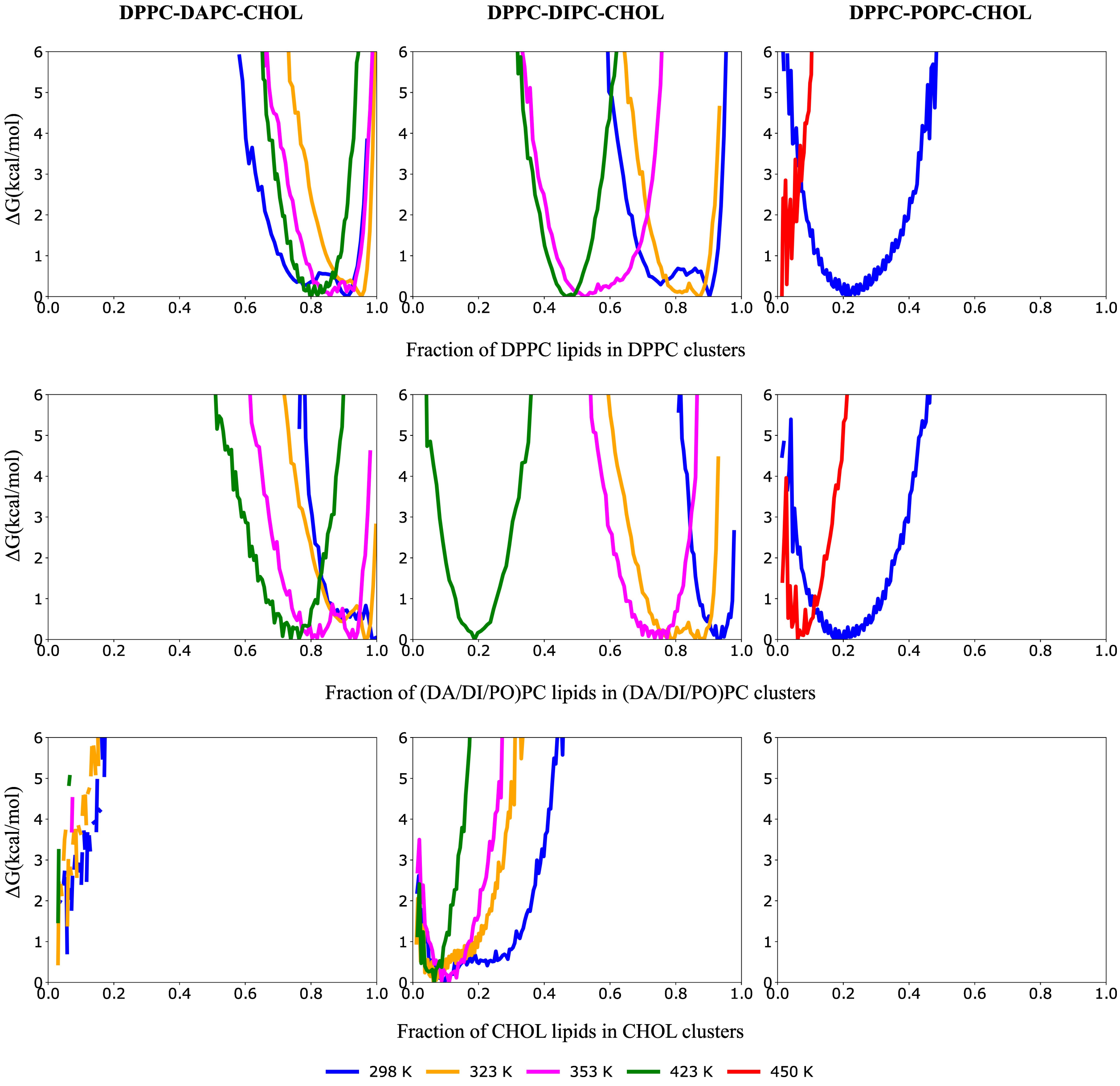


Figure S3: Species-wise FLC ΔG profile for each system as a function of temperatures. Each column represents each lipid system under study. The rows represent the saturated, unsaturated and cholesterol species within each system respectively

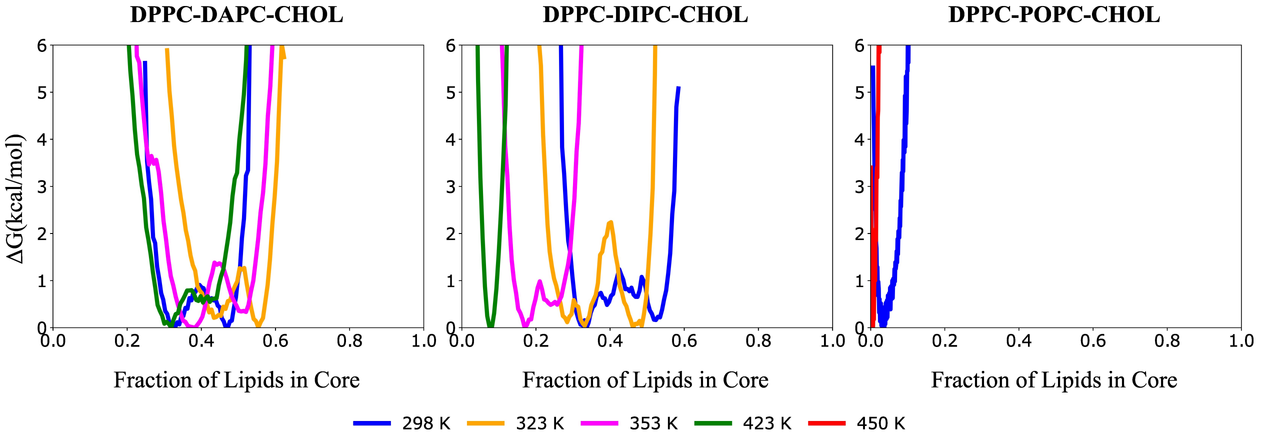


Figure S4: Fraction of Lipids in Core ΔG profile for each system as a function of temperatures. Each column represents each lipid system under study. The rows represent the total, saturated, unsaturated and cholesterol species within each system respectively

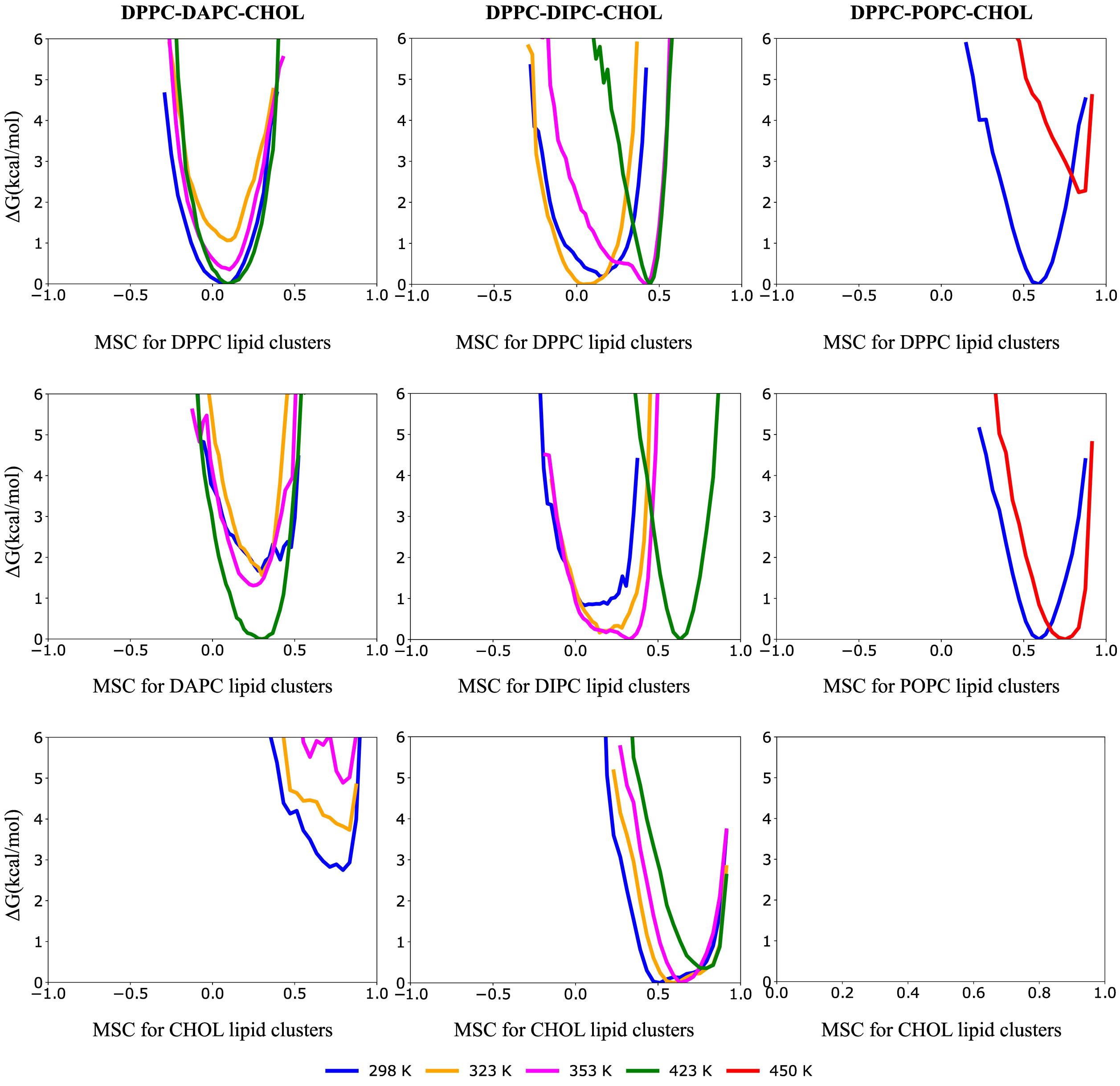


Figure S5: Mean MSC vs. ΔG profile for each system as a function of temperatures. Each column represents each lipid system under study. The rows represent the saturated, unsaturated and cholesterol species within each system respectively

### 5 Role of cholesterol

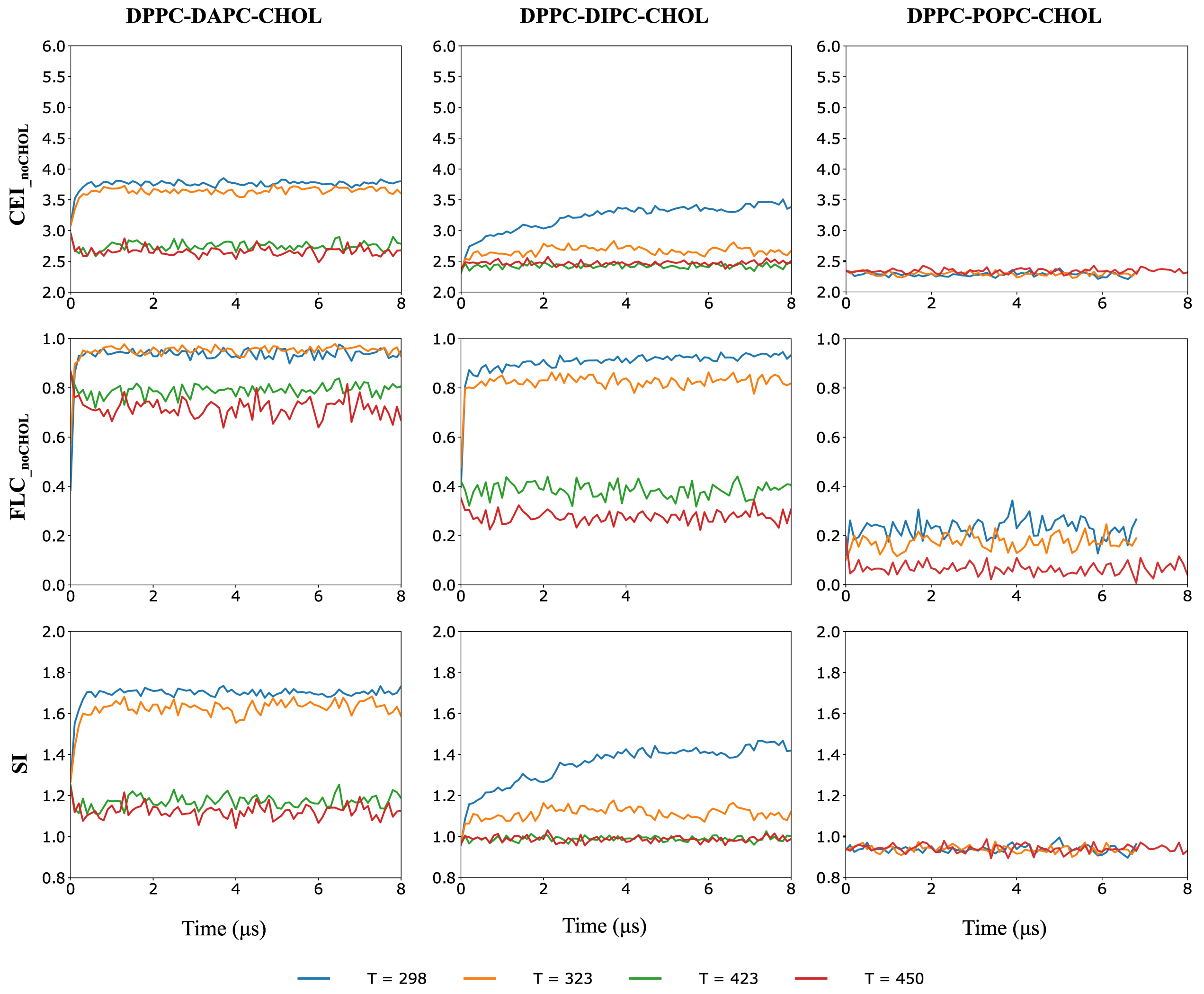


Figure S6: Complementary plots to the Fig 3 in Main Article that illustrates the role of cholesterol in CV calculation. Each column represents each lipid system under study. The first two rows represents CEI and FLC without the cholesterol contribution. While the third row represents SI with cholesterol contribution

#### 6 Convergence of free energy curves across replicas

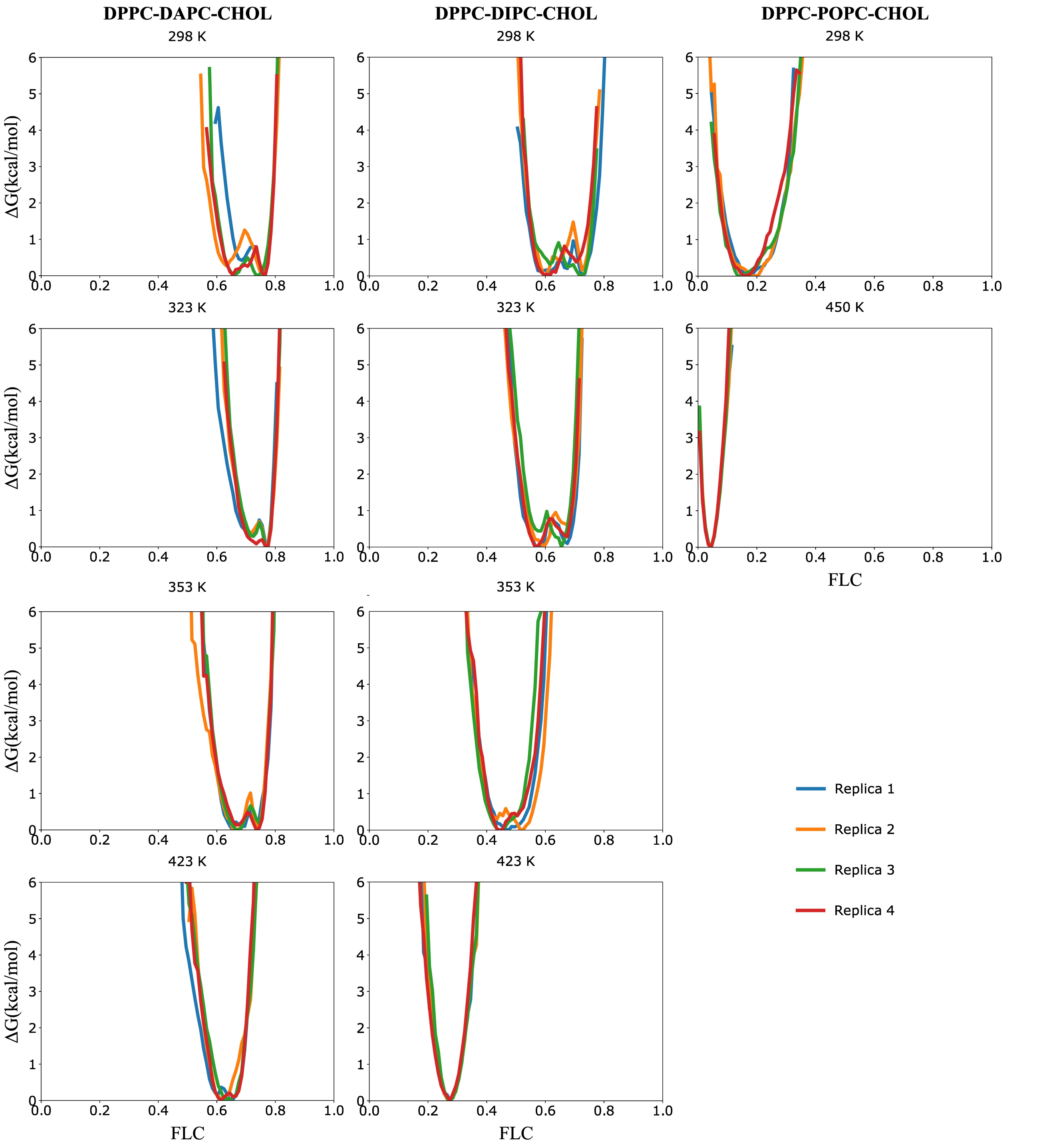


Figure S7: The convergence between four replicas of the system at different temperatures. Each column represents each lipid system under study.

### 7 $\Delta\Delta G_{\text{sep}}$ estimation

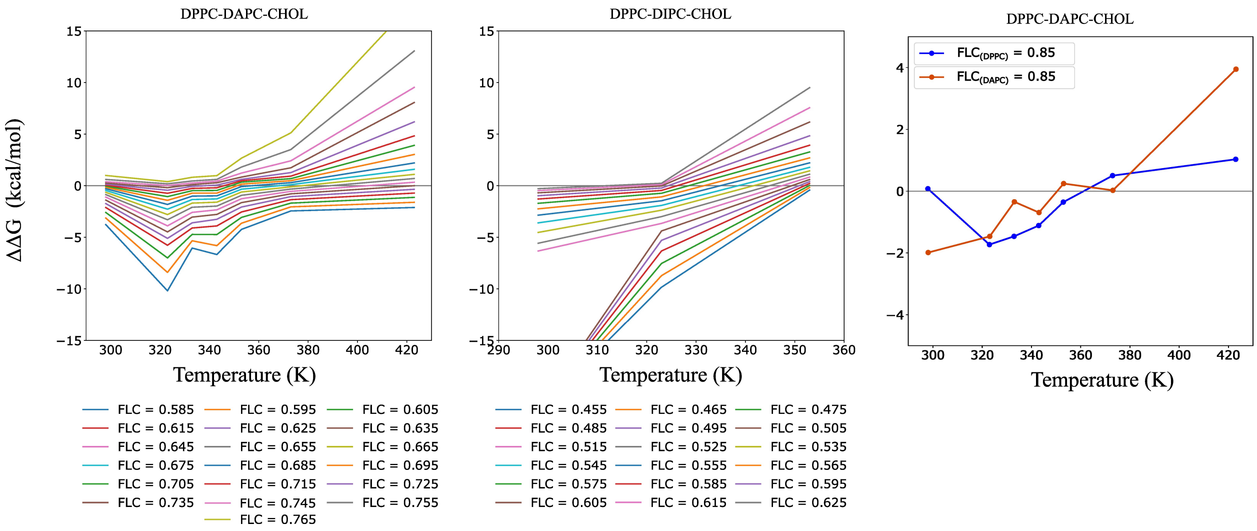


Figure S8: Sensitivity of $\Delta\Delta G_{\text{sep}}$ estimation to the choice of FLC cutoff for each system.

[7] Mark Abraham, Andrey Alekseenko, Cathrine Bergh, Christian Blau, Eliane Briand, Mahesh Doijade, Stefan Fleischmann, Vytautas Gapsys, Gaurav Garg, Sergey Gorelov, Gilles Gouaillardet, Alan Gray, M. Eric Irrgang, Farzaneh Jalalypour, Joe Jordan, Christoph Junghans, Prashanth Kanduri, Sebastian Keller, Carsten Kutzner, Justin A. Lemkul, Magnus Lundborg, Pascal Merz, Vedran Miletić, Dmitry Morozov, Szilárd Páll, Roland Schulz, Michael Shirts, Alexey Shvetsov, Bálint Soproni, David van der Spoel, Philip Turner, Carsten Uphoff, Alessandra Villa, Sebastian Wingbermühle, Artem Zhmurov, Paul Bauer, Berk Hess, and Erik Lindahl. GROMACS 2023 Manual. feb 2023.
